# Proteomics of human cancer-associated T cells identifies regulators of T cell functionality

**DOI:** 10.64898/2026.08.18.745433

**Authors:** Kaspar Bresser, Žan Hozjan, Nila H Servaas, Suzan Stelloo, Cornelia G Spruijt, Maïa Nestor Martin, Aurélie Guislain, Nandhini Kanagasabesan, Stijn Kneefel, Arie Johan Hoogendijk, Carmen van der Zwaan, Živa Moravec, Rhianne Voogd, Marja Nieuwland, Jessica Sieljes, Robert van Es, Kim Monkhorst, Koen Hartemink, Willemijn SME Theelen, Wouter Scheper, Michiel Vermeulen, Monika C Wolkers

**Author notes:** Corresponding author: Monika C Wolkers. Authors contributed equally (Kaspar Bresser, Žan Hozjan).

## Abstract

CD8⁺ T cells in solid cancers progressively lose anti-tumor activity, yet the cell-intrinsic mechanisms driving this loss of function remain incompletely defined. Here, we performed matched proteomic and transcriptomic profiling of dysfunctional and bystander CD8⁺ tumor-infiltrating T cells isolated from primary tumors of treatment-naïve non-small cell lung cancer patients. Proteomic analysis revealed widespread discordance with mRNA expression, with 8% of all quantified proteins displaying differential expression exclusively at the protein level. Genetic perturbation of such differentially expressed proteins identified the chromatin remodeler CHD4 and fatty acid synthase (FASN) as cell-intrinsic regulators of T cell function. CHD4 deletion resulted in altered gene-regulatory networks that promoted effector differentiation and enhanced cytokine production. In contrast, FASN deletion preserved mitochondrial fitness and sustained T cell functionality under chronic T cell receptor stimulation. Together, these findings demonstrate that proteomic profiling uncovers regulators of T cell functionality that are not apparent from transcriptomic analyses alone, highlighting an additional layer of regulatory control.

**One Sentence Summary:** Integrated multi-omic profiling of human tumor-infiltrating T cells reveals cell-intrinsic regulators of T cell dysfunction that are missed by transcriptomic analyses alone.

## Introduction

CD8⁺ T cells undergo extensive molecular and functional reprogramming during antigen encounter, differentiating into cytotoxic effector cells capable of eliminating infected and malignant cells(*1*, *2*). Effector T cells achieve this through the release of cytotoxic granules, together with the production of pro-inflammatory cytokines, including interferon-γ (IFNγ) and tumor necrosis factor (TNF) following activation through the T cell receptor (TCR). However, in persistent antigen stimulation, such as cancer, this differentiation trajectory is disrupted, and CD8⁺ T cells instead acquire a dysfunctional state(*1*, *3*). These cells are characterized by sustained expression of inhibitory receptors such as PD-1, CD39 and LAG-3 together with impaired cytokine production, resulting in a profound loss of effector function(*1*, *4*, *5*). This dysfunctional state has been proposed to limit immunopathology during chronic immune responses, but in doing so it enables tumor progression. Moreover, T cell dysfunction is accompanied by widespread remodeling of the genetic enhancer landscape and transcription factor binding, which leads to a distinct and stable epigenetic state(*5*, *6*). Mitochondrial defects further stabilize the dysfunctional T cell state by compromising cellular metabolic fitness(*7*, *8*).

Single-cell RNA sequencing has revolutionized the study of cancer-associated T cells, revealing heterogeneous transcriptional states and diverse levels of dysfunction shaped by the tumor microenvironment(*4*, *9*, *10*). Such studies identified central regulators of T cell dysfunction, including the transcription factors *TOX*(*11*, *12*) and *RBPJ*(*13*). While these scRNAseq studies were instrumental in deepening our understanding of T cell dysfunction, mRNA abundance does not always correlate with protein expression or functional capacity(*14–17*). In T cells, the discrepancies between transcriptomic and proteomic measurements are increasingly recognized to be shaped by post-transcriptional regulation, including mRNA stability, translation efficiency, and localization(*18–20*). This discrepancy limits the extent to which transcriptional data alone can describe T cell states.

Recent studies have begun to explore the proteomic landscape of tumor-infiltrating CD8⁺ T cells(*7*, *21*–*23*). For example, proteomic analyses of cancer-associated T cells revealed a pathological protein stress response in dysfunctional T cells, characterized by protein aggregation and selective chaperone imbalance, which together compromise effector function(*22*). Proteomic studies have additionally identified novel therapeutic targets for potentially reinvigorating dysfunctional T cells(*21*, *23*). Integrating proteomic and transcriptomic analyses may therefore uncover additional regulators of T cell function and differentiation that are missed by mRNA-based approaches alone. Furthermore, proteomic analyses of human cancer-associated T cells have been performed on bulk tumor-infiltrating T cell populations. However, the large influx of bystander T cells(*24*) (e.g., from the tumor vasculature or recruited through inflammatory cues) may dilute subset-specific molecular signatures of dysfunctional CD8⁺ T cells.

To address these limitations, we performed proteomic profiling of FACS-sorted human CD8⁺ T cell subsets isolated from tumors and adjacent lung tissue of non-small lung cell cancer (NSCLC) patients. By separating dysfunctional and bystander tumor-infiltrating T cell populations, we identified protein expression profiles and pathways that define distinct cancer-associated T cell states. Comparative analysis of transcriptomics and proteomics revealed a widespread mismatch between mRNA and protein abundance, uncovering a large set of differentially expressed proteins that were not apparent from transcriptomic analyses alone. Functional validation of these differentially expressed proteins identified FASN and CHD4 as key regulators of T cell function. We found that CHD4-dependent chromatin remodeling shapes transcriptional programs linked to effector T cell differentiation. In contrast, FASN contributes to chronic TCR stimulation induced metabolic stress and mitochondrial defects that promote T cell dysfunction. Notably, depletion of these factors enhanced T cell effector function. Together, our findings demonstrate that proteomic profiling exposes an additional layer of T cell regulatory biology that remains obscured at the transcriptomic level and provides a framework for identifying functionally relevant regulators of T cell state.

## Results

### CD103 and CD39 mark dysfunctional T cells in NSCLC lesions

To study the proteome of tumor-infiltrating dysfunctional T cells, we used enzymatically digested, cryopreserved tumor digests from NSCLC patients, together with matched digests from distal lung tissue as control(*25*, *26*). We focused our study on early-stage treatment-naïve NSCLC patients to avoid therapy-induced confounding effects **(Figure 1a**). To identify suitable surface markers that separate dysfunctional from bystander T cells, we evaluated previously implicated proteins, including CD39, PD- 1, and CD103(*27*), using single-cell RNA sequencing (scRNAseq) with TCR profiling and antibody- derived tagging of tumor-derived CD8^+^ T cells (**Figure 1b**).

**Figure 1.**
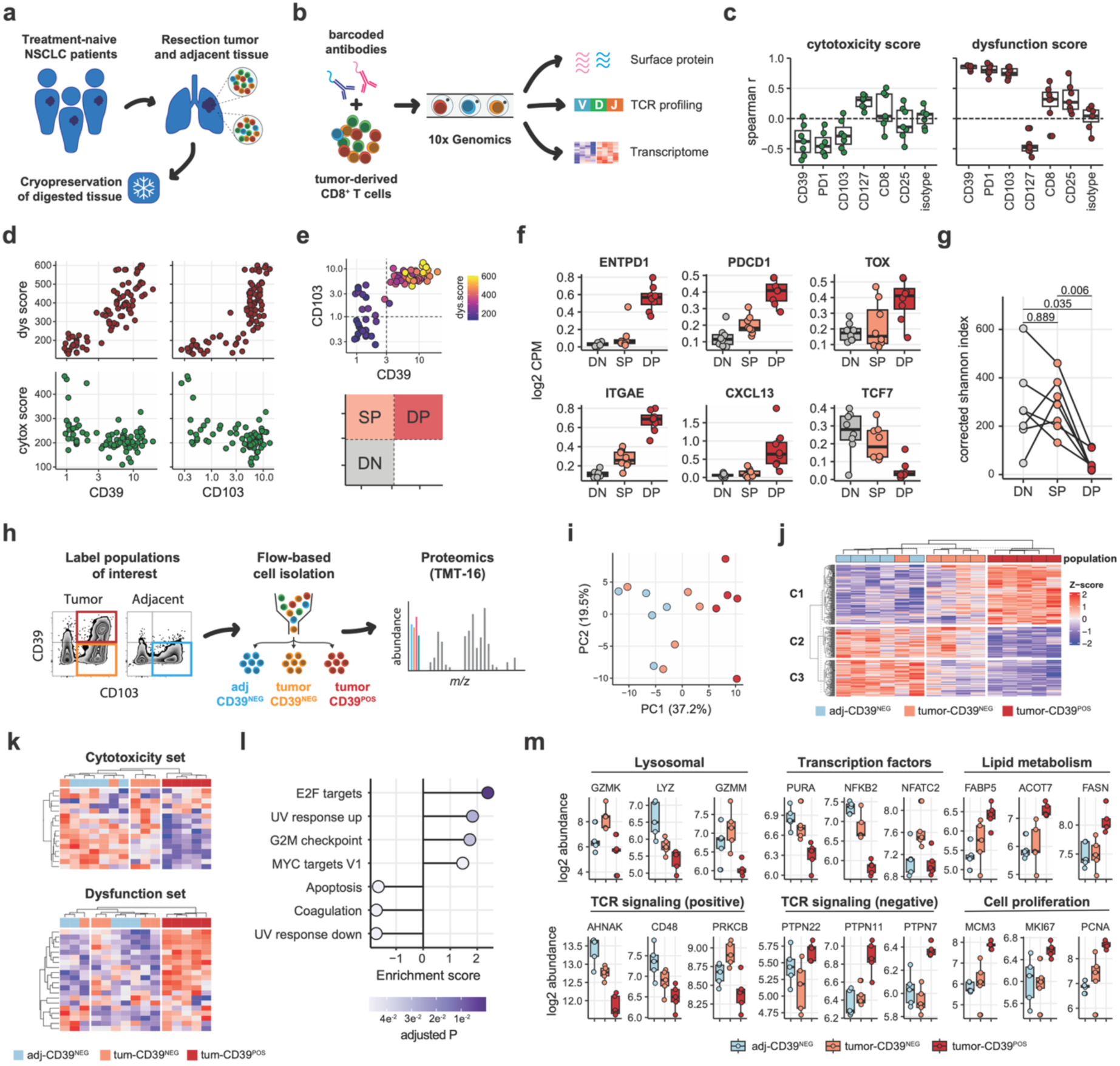
Proteomic assessment of human cancer-associated CD8^+^ T cells. (**a**) Primary material obtained from non-small cell lung cancer (NSCLC) patients. (**b**) Single-cell RNA sequencing (scRNAseq) coupled with TCR profiling and barcode-labeled antibodies to identify dysfunctional T cells. (**c**) Spearman correlations between transcriptome-based scores calculated from T cell dysfunction and cytotoxicity gene-sets(*29*) and quantified binding of antibodies directed against indicated proteins. (**d-e**) Relationship between dysfunction and cytotoxicity scores and surface expression of indicated proteins. Dots indicate MetaCell clusters (see methods). (**f**) Pseudobulk read counts of indicated genes calculated for the three populations indicated in panel (**e**). Dots indicate individual patients. (**g**) Shannon diversity index of T-cell receptor CDR3 clones. Index is corrected for sample size bias. Dots indicate individual patients. (**h**) Experimental setup for proteomic analysis of cancer-associated T cell populations. (**i**) Principal component analysis of the top 500 variable proteins. Dots indicate individual samples. (**j**) Hierarchical clustering of all differentially expressed proteins across assessed T cell populations. Marker proteins for each row-cluster are displayed in **Supplementary Figure 2c**. (**k**) Hierarchical clustering of proteins associated with T cell cytotoxicity and dysfunction(*29*). (**l**) Gene-set enrichment analysis (Hallmark pathways, Msigdb). (**m**) Selected proteins involved in T cell function displaying significant differential expression across the assessed T cell populations. Plots show protein abundance (LFQ). Dots indicate individual samples. scRNAseq (n=7 patients) and proteomics (n=5 patients) data were obtained from a single cohort of treatment-naïve NSCLC patients. Boxplots (**f, m**) show the median and 25th/75th percentiles, whiskers extend to 1.5× the interquartile range. P values were determined by multiple paired two-sided Student’s t-tests (**g**) with Holm–Bonferroni correction for multiple testing.

Grouping T cells into representative “MetaCells”(*28*) (**Supplementary Figure 1**) revealed that CD39 and CD103 expression levels strongly associated with core dysfunction genes and inversely correlated with a T cell cytotoxicity gene signature (**Figure 1c-d, Supplementary Table 1**). Furthermore, pseudo-bulking CD8^+^ T cells based on CD39/CD103 surface expression (**Figure 1e, Supplementary Figure 1c**) showed that CD103^hI^CD39^hI^ CD8^+^ T cells expressed high levels of dysfunction-associated genes, including *PDCD1* (encoding PD-1), *CXCL13*, and *TOX* (**Figure 1f**). This T cell population was also enriched for pathways related to T cell dysfunction in chronic infections (**Supplementary Figure 1d**). Moreover, the CD103^hI^CD39^hI^ population exhibited the lowest TCR diversity (**Figure 1g**), indicating antigen-driven clonal expansion—a hallmark of T cell dysfunction(*29*). Together, these data indicate that the CD103^hI^CD39^hI^ subset represents the *de facto* dysfunctional T cell population in this cohort.

### Mass-spectrometric analysis of cancer-associated T cells

Based on these data, we isolated CD8^+^ T cell populations by fluorescence-activated cell sorting (FACS) from 5 NSCLC tumors, dividing them into “bystander” CD103⁺CD39⁻ single-positive (tumor-CD39^NEG^) T cells and “dysfunctional” CD103⁺CD39⁺ double-positive (tumor-CD39^POS^) T cells (**Figure 1h, Supplementary Figure 2a**). As control, we isolated “tissue-patrolling” CD103⁺CD39⁻ CD8^+^ T cells from tumor-adjacent tissue (adj-CD39^NEG^). Per population, 50,000–150,000 T cells were collected (**Supplementary Figure 2b**). The limited biomass of T cells(*30*) poses a challenge for protein quantification by LC-MS. To achieve comparative proteomic analysis at sufficient depth, samples were individually lysed and multiplexed using isobaric labeling (TMT-16). This strategy enabled the quantification of 1,827 proteins across all T cell populations (**Supplementary Table 2**).

Principle component analysis revealed that adj-CD39^NEG^ and tumor-CD39^NEG^ CD8^+^ T cells from all NSCLC patients clustered closely together, whereas tumor-CD39^POS^ CD8^+^ T cells formed a more distinct group (**Figure 1i**). Interestingly, unsupervised clustering revealed that tumor-CD39^NEG^ T cells primarily displayed an intermediate protein expression profile (**Figure 1j, Supplementary Figure 2c**), possibly reflecting their adaptation to the tumor microenvironment. As expected, tumor-CD39^POS^ T cells expressed higher levels of proteins that mark T cell dysfunction and reduced levels of proteins associated with T cell cytotoxicity (**Figure 1k, Supplementary Table 1**). In addition, tumor-CD39^POS^ T cells were enriched for pathways implicated in T cell dysfunction, including proliferation, DNA damage responses, and apoptosis (**Figure 1l-m, Supplementary Table 3**). Consistently, tumor-CD39^POS^ T cells exhibited decreased expression of positive regulators of TCR signaling (e.g., AHNAK, PRKCB) and increased expression of negative regulators, including the tyrosine phosphatases PTPN7 and PTPN11 (**Figure 1m**). Key transcriptional regulators of T cell function, including PURA, NFKB2, and NFATC2, were also downregulated (**Figure 1m**). In line with recent studies implicating altered lipid metabolism in T cell dysfunction(*31*, *32*), expression of lipid metabolism related proteins FABP5 and FASN was upregulated in tumor-CD39^POS^ CD8^+^ T cells. To assess whether these observations extended beyond CD8⁺ T cells, we performed analogous proteomic profiling on matched CD4⁺ T cell populations isolated from the same NSCLC samples based on the expression of CD69 and CD39(*33*) (**Supplementary Figure 2a**). Although CD4⁺ T cell subpopulations displayed less distinct proteomic clustering, they exhibited similar proteomic features associated with chronic activation and altered T cell function (**Supplementary Figure 2d-g, Supplementary Table 4**). Together, these data demonstrate that multiplexed LC-MS enables proteomic interrogation of distinct T cell subpopulations within the tumor microenvironment.

### Proteomics and mRNA sequencing capture different components of T cell dysfunction

We next performed matched bulk mRNA sequencing on the same sorted CD8^+^ T cell populations used for proteomic analysis to assess the relationship between protein and mRNA abundance (**Supplementary Figure 3**). Consistent with previous studies in different T cell subsets(*19*, *34*, *35*), comparison of differential expression at the protein and mRNA levels revealed only a limited positive correlation (Spearman r = 0.30–0.38; **Figure 2a**). Strikingly, only 11–35% of differentially expressed proteins were also differentially expressed at the mRNA level (**Figure 2b-c, Supplementary Table 5**), demonstrating a substantial discordance between transcriptomic and proteomic measurements. As a result, proteomic and transcriptomic analyses identified distinct pathways relevant to T cell biology. While both analyses captured differential expression of E2F-targets and interferon-responsive genes, mRNA sequencing uniquely identified alterations in glycolysis and mTORC1 signaling in tumor-CD39^POS^ CD8^+^ T cells (**Figure 2d, Supplementary Table 6**). In contrast, expression level differences of MYC targets, the DNA damage response, and cytokine signaling were only apparent at the protein level across the different comparisons (**Figure 2d**). Notably, concordance between mRNA and protein measurements was not uniform within gene classes or pathways. Both concordant and discordant protein–mRNA relationships were observed among genes involved in key aspects of T cell function, such as secretory lysosomes, TCR signaling, and transcriptional regulation. Concordant regulation was observed for GZMA, GZMK, AHNAK, NFATC2, and STAT4, whereas NFKB2, SUZ12, LYZ, and CD48 showed marked discordance between protein and mRNA abundance (**Figure 2e**). In summary, this comparative analysis highlights that although mRNA sequencing and proteomics display some level of redundancy, each methodology captures distinct and complementary components of T cell dysfunction that would be incompletely defined when assessed in isolation.

**Figure 2.**
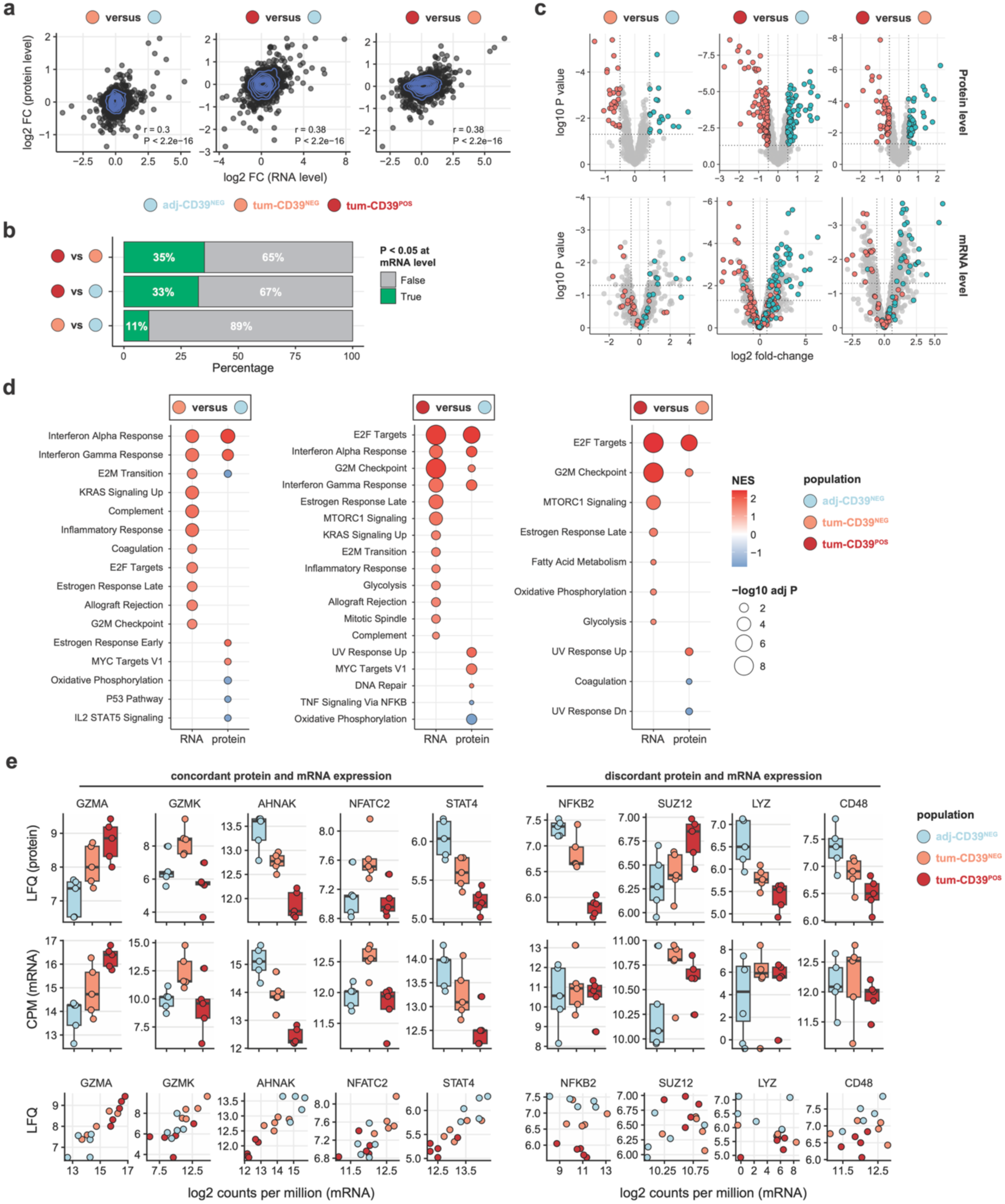
Proteomics and mRNA sequencing capture different components of T cell dysfunction. (**a**) Association between fold changes observed at the RNA and protein level, across the assessed CD8^+^ T cell populations. (**b**) Percentage of differentially expressed proteins, stratified by whether they are also significantly differentially expressed at the RNA level. (**c**) Differential expression analysis of LC-MS (top) and mRNAseq (bottom) data for the indicated comparisons. Proteins with P values < 0.05 in the LC-MS dataset are highlighted in both graphs. (**d**) Gene-set enrichment analysis (Hallmark pathways, Msigdb) on mRNA or protein level for indicated comparisons. (**e**) Protein and mRNA abundance of selected genes displaying discordance, with correlation plots showing the relationship of abundances across T cell populations. Matched mRNAseq and proteomics data were obtained from 5 treatment-naïve NSCLC patients. Boxplots (**e**) show the median and 25th/75th percentiles, whiskers extend to 1.5× the interquartile range.

### Genetic perturbation of FASN and CHD4 influences T cell state

The widespread discordance between mRNA and protein expression raised the possibility that functionally important regulators of T cell function may remain undetected by transcriptomic analyses alone. Therefore, we examined whether proteins identified exclusively at the protein level intrinsically regulate T cell differentiation and function. To this end, we selected nine discordant proteins (**Supplementary Figure 4a**): four putative negative regulators (enriched in tumor-CD39^POS^ T cells; TPP2, DDX3, CHD4, FASN) and five putative positive regulators of T cell function (depleted in tumor-CD39^POS^ T cells; HMGN4, PTGES2, CRYZ, PRCP, APEX1). None of these nine proteins displayed differential mRNA expression between tumor-CD39^POS^ and tumor-CD39^NEG^ T cells across five external scRNAseq datasets (**Supplementary Figure 4b-c**). In addition to these discordant proteins, we included five proteins (PTPN22, ACAT2, CARPIN1, STAT3, ACLY) that displayed strong concordance between mRNA and protein expression. To test the involvement of these proteins in T cell biology, we performed CRISPR- Cas9-mediated knock-out of the selected genes on activated blood-derived naïve CD8⁺ T cells (**Supplementary Figure 4d**). Expression of 12 markers of T cell state was quantified by flow cytometry 7 days post-activation (5 days after gene-editing) and compared to T cells that were transfected with a non-targeting (NT) Cas9 complex (**Supplementary Table 7, Supplementary Data 1**). Although genetic perturbation of most targets induced some degree of phenotypic change, deletion of the chromatin remodeler CHD4 and fatty acid synthase FASN produced the most pronounced alterations. This included altered protein expression of key markers such as CCR7, CD39, LAG3, and T-bet (**Supplementary Figure 4e-f**). CHD4 and FASN are putative negative regulators of T cell function, as their protein levels were increased in both dysfunctional CD8^+^ and CD4^+^ T cells (**Supplementary Figure 2g**). Given their roles in epigenetic regulation (CHD4) and cellular metabolism (FASN)—two major processes disrupted in dysfunctional CD8^+^ T cells—we focused our subsequent analyses on these two proteins.

### CHD4 restrains T cell effector differentiation and functional maturation

CHD4 is an integral subunit of the NuRD chromatin remodeling complex, a key mediator of epigenetic lineage repression during cellular differentiation(*36*, *37*), and of the chromatin looping complex ChAHP(*38*). CHD4 has additionally been implicated in the thymic development of CD3^+^ T cells(*39*) and identified as a regulator of T helper 1 cell polarization(*40*). Therefore, we hypothesized that CHD4 may play a role in the epigenetic regulation of CD8^+^ T cell state. To test this, we performed an extensive phenotypic analysis of CHD4-knockout (CHD4-KO) T cells at 7 days post-activation (**Figure 3a, Supplementary Figure 5a**). CHD4-KO T cells showed substantially reduced expression of CD27 and CCR7 compared to control T cells (**Figure 3b**), indicative of enhanced effector differentiation. Simultaneously, the T cell activation-associated receptors CD39, LAG3, and PD-1 were elevated in CHD4-KO T cells (**Supplementary Figure 5b**). Consistent with the observed effector phenotype, CHD4- KO T cells exhibited increased expression of the effector-lineage transcription factor T-bet (**Figure 3c**). The expression of additional effector-associated transcription factors (TOX and BLIMP-1; **Figure 3c, Supplementary Figure 5c**) was also upregulated, supporting a role for CHD4 in controlling the transcriptional circuitry linked to effector function(*41*, *42*). Interestingly, TCF1 expression was also maintained to a higher extent in CHD4-KO T cells (**Figure 3c, Supplementary Figure 5c**), which is normally repressed during antigen-driven proliferation and effector maturation(*43*). This finding suggests that CHD4 loss promotes non-canonical effector T cell differentiation characterized by partial retention of TCF1 expression.

**Figure 3.**
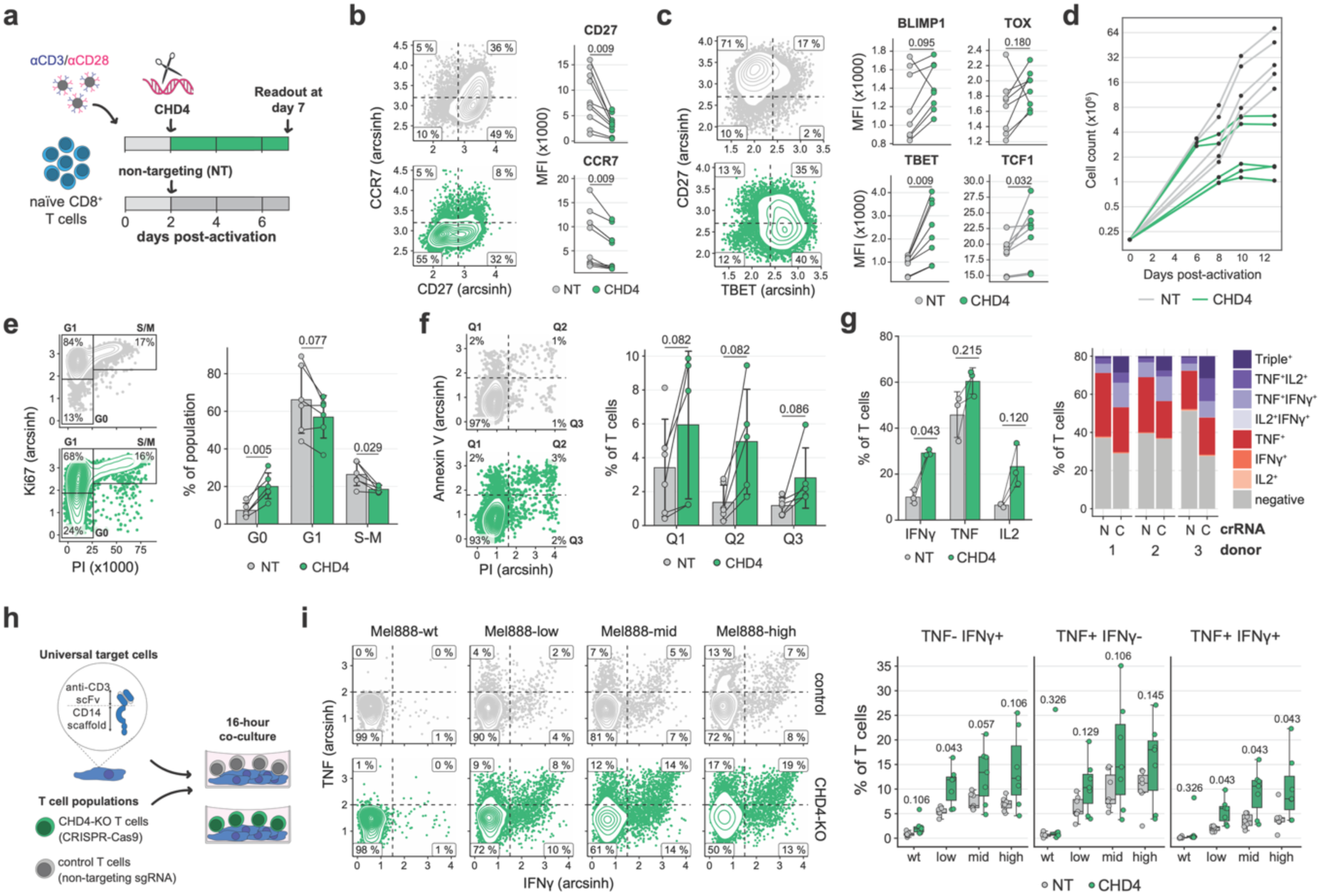
CHD4 deletion promotes T cell effector differentiation and functional maturation. Human naïve CD8^+^ T cells were activated with anti-CD3/CD28 dynabeads, nucleofected with *CHD4* or non-targeting Cas9–guide RNA complexes (NT) at day 2 post-activation and analyzed at day 7 post-activation. (**a**) Experimental timeline. (**b-c**) Protein expression levels for indicated surface proteins (**b**) and transcription factors (**c**), measured by flow cytometry. Representative density plots (left) and summarizing strip charts (right) are shown. Lines connect samples obtained from individual donors (n = 8-10). (**d**) Cell expansion curves. Lines indicate samples obtained from individual donors (n = 5). (**e-f**) Cell cycle analysis (**e**) and Annexin-V and propidium iodide staining (**f**). Representative density plots (left) and summarizing bar plots (right) are shown. Dots indicate samples obtained from individual donors (n = 6), bars indicate group means, error bars indicate standard deviation. (**g**) Intracellular cytokine staining of indicated cytokines after a 3-hour re-stimulation with anti-CD3/CD28 antibodies or PMA/ionomycin in the presence of brefeldin A. Dots indicate samples obtained from individual donors (n = 3), bars indicate group means, error bars indicate standard deviation. Polyfunctionality analysis showing the percentage of cells producing TNF, IFNγ, and IL-2, including single-, double-, and triple-cytokine–producing T cells. (**h-i**) Co-culture assay of T cells with Mel888 cells that were engineered to express a membrane-tethered single-chain anti-CD3 antibody at different expression levels. (**h**) Experiment setup. (**i**) Intracellular cytokine staining of indicated cytokines after 16-hours of co-culture (E:T ratio 1:1) with indicated Mel888 lines in the presence of Brefeldin A (added after the first 20 minutes of the culture). Representative density plots (left) and summarizing bar plots (right) are shown. Dots indicate samples obtained from individual donors (n = 7). Displayed graphs are representative of 2-4 independent experiments, and contain data from 1 experiment (**d, g**) or compiled from 2 (**e, f**) or 3 (**b, c**) experiments. Boxplots (**i**) show the median and 25th/75th percentiles, whiskers extend to the minimum/maximum values. P values in all panels were determined by multiple paired two-sided Student’s t-tests with Benjamini–Hochberg FDR correction for multiple testing.

Progressive effector T cell differentiation is typified by reduced proliferative potential(*44*, *45*), increased sensitivity to apoptosis(*46*), and alterations in functional capacity (e.g., cytokine production profiles)(*2*, *47*). Consistent with the acquisition of an effector-associated state, CHD4-KO cells displayed reduced expansion capacity (**Figure 3d**) and proliferative activity (**Figure 3e**). In addition, we observed a modest increase in the percentage of early apoptotic, late apoptotic and dead cells in CHD4-KO populations (**Figure 3f**), showing that CHD4 loss limits proliferative activity and increases sensitivity to apoptosis. We next tested for alterations in the functional capacity, by re-stimulating CHD4-KO CD8^+^ T cells with anti-CD3/CD28 antibodies or PMA/Ionomycin. CHD4-KO T cells produced significantly higher amounts of the pro-inflammatory cytokines IL-2, IFNγ, and TNF compared to control T cells (**Figure 3g, Supplementary Figure 5**). Notably, CHD4-KO populations exhibited increased frequencies of double and triple positive cytokine producing T cells, indicating enhanced polyfunctionality (**Figure 3g**). To further corroborate these findings in a cellular context, we co-cultured CHD4-KO and control CD8^+^ T cells for 16 hours with engineered MEL888 cells expressing graded levels of membrane-tethered anti-CD3 antibodies (**Supplementary Figure 5**), mimicking different levels of antigen-dependent TCR crosslinking(*48*) (**Figure 3h**). CHD4-KO T cell populations consistently displayed higher frequencies of IFNγ and TNF producing cells (**Figure 3i, Supplementary Figure 5**). Additionally, the percentage of IFNγ and TNF double-producing T cells was significantly increased, indicating enhanced sensitivity of CHD4-deficient T cells to TCR engagement. Together, these data show that CHD4 restrains effector-associated differentiation and limits the functional responsiveness of activated CD8⁺ T cells.

### CHD4 drives effector-associated transcriptional programs through chromatin remodeling

We next sought to understand how CHD4 was driving the effector T cell state. Co-immunoprecipitation (Co-IP) of CHD4 confirmed its interaction with both the NuRD and the ChAHP complex in activated CD8^+^ T cells (day 7 post-activation; **Figure 4a, Supplementary Table 8**). Both the NuRD complex, which couples CHD4 with histone deacetylases HDAC1/2, and the related Polycomb repressive complex 2 have been previously implicated in the epigenetic regulation of T cell effector function(*49–51*). As CHD4 is part of the core NuRD complex, we hypothesized that deletion of CHD4 may destabilize this complex, leading to increased gene accessibility at effector-associated loci. However, deletion of CHD4 in CD8^+^ T cells did not affect the cellular levels of HDAC1 or HDAC2 (**Figure 4b**). Importantly, Co-IP of HDAC1 efficiently recovered all components of the NuRD complex in both CHD4-KO (**Figure 4c, Supplementary Table 9**) and control T cells (**Supplementary Figure 6a-b**), indicating that the NuRD complex remained intact upon CHD4 deletion. Intriguingly, the neuronal-associated paralog of CHD4, CHD5, was found to interact with HDAC1 in CHD4-KO T cells (**Figure 4d**), suggesting that CHD5 may partially compensate for CHD4 loss within the NuRD complex.

**Figure 4.**
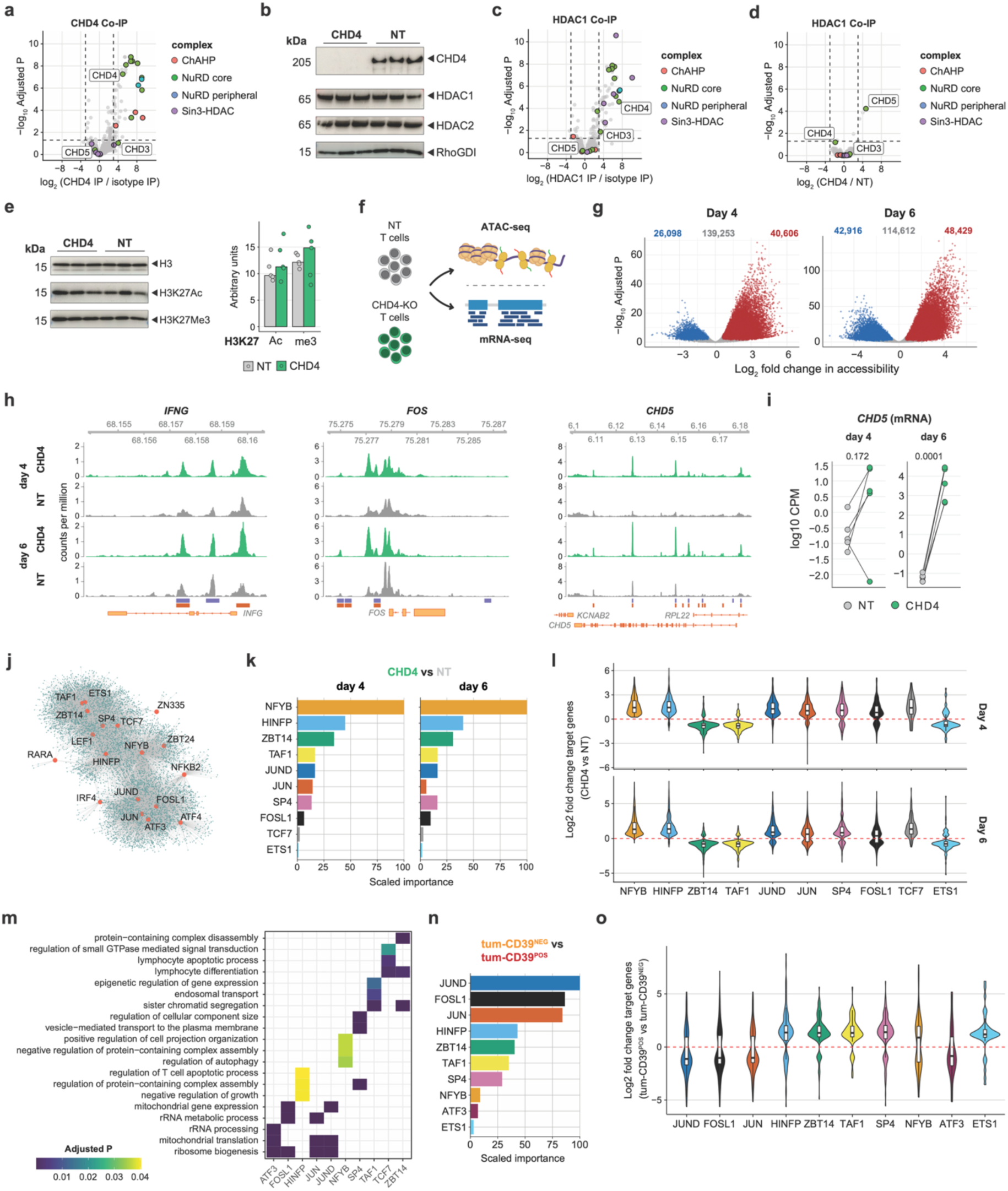
CHD4 is involved in the epigenetic repression of effector T cell differentiation. (**a**) Co-immunoprecipitation on T cell lysates (day 7 post-activation) using an anti-CHD4 antibody and isotype control (n = 3). (**b**) Western blots for indicated known CHD4 interactors. (**c**) Co-immunoprecipitation on T cell lysates using an anti-HDAC1 antibody and isotype control (n = 3). (**d**) Co-immunoprecipitation on CHD4-proficient and - deficient T cell lysates using an anti-HDAC1 antibody and isotype control (n =3). (**e**) Western blots for indicated histone modifications. Dots indicate individual donors (n = 5). (**f**) mRNA-seq and ATAC-seq was performed on CHD4-proficient and -deficient T cells (from matched donors) at day 4 and day 6 post activation. (**g**) Differential accessibility of regions identified by ATAC-seq. (**h**) Chromatin accessibility profiles measured by ATAC-seq for *IFNG, FOS* and *CHD5*. Blocks below tracks indicate significant (adj P < 0.05) differentially accessible regions on da 4 (purple) and day 6 (orange). (**i**) mRNA expression levels of the *CHD5* gene. Dots indicate individual donors (n = 4-5). (**j**) Enhancer-based gene regulatory network (eGRN) based on covariation of chromatin accessibility and RNA-seq data across samples. Red dots indicate transcription factors, lines represent identified enhancer regions, turquoise dots indicate target genes. (**k**) eGRN-based transcription factor importance for predicting differential gene expression in CHD4-KO versus NT T cells. Results for day 4 (top) and day 6 (bottom) post-activation (day 2 and 4 post gene-editing) are displayed. (**l**) Differential expression of target genes of the indicated transcription factors. Results for day 4 (top) and day 6 (bottom) are displayed. (**m**) Pathway overrepresentation analysis within the target genes for indicated transcription factors. (**n**) eGRN-based transcription factor importance for predicting differential gene expression in t-DP T cells versus t-SP T cells. (**o**) Differential expression (t-DP T cells versus t-SP T cells) of target genes of indicated transcription factors. Data are representative of 2 independent experiments (**a-e**) or obtained in a single experiment (**f-o**). Boxplots (**l, o**) show median and 25th/75th percentiles, whiskers extend to 1.5× the interquartile range. P values were determined by empirical Bayes-moderated t-tests with Benjamini–Hochberg correction for multiple testing (**a, c, d**) or multiple paired two-sided Student’s t-tests with Holm–Bonferroni correction for multiple testing (**i**).

To assess whether CHD4 deletion affects histone modifications, we measured global H3K9Ac, H3K27Ac, and H3K27Me levels by Western blot. No differences were observed between CHD4-KO and control CD8^+^ T cells, indicating that CHD4 loss does not alter the total histone modification levels (**Figure 4e**). We next investigated if CHD4 regulates gene-specific chromatin accessibility and downstream transcriptional programs by performing matched mRNA-seq and ATAC-seq on CHD4-KO and control CD8^+^ T cells (**Figure 4f, Supplementary Figure 6c-d, Supplementary Table 10**). To capture both early and more sustained effects of CHD4 deficiency, samples were collected at days 4 and 6 post- activation (i.e., 2 and 4 days after gene-editing). ATAC-seq identified 205,957 consensus regions across all samples (**Figure 4g**). Data quality was supported by a high signal-to-noise ratio and strong enrichment at transcription start sites (**Supplementary Figure 6e-h**). Consistent with the role of CHD4 in maintaining repressive chromatin states, differential accessibility analysis revealed predominantly increased chromatin accessibility in CHD4-KO T cells at day 4 (20% increased versus 13% decreased regions). By day 6, gains and losses in chromatin accessibility occurred at similar frequencies (24% increased versus 21% decreased regions; **Figure 4g, Supplementary Table 11**). In line with our findings linking CHD4 to effector maturation (Figure 3), CHD4-KO T cells displayed enhanced accessibility at the loci of the pro-inflammatory cytokine *IFNG* and the transcription factor *FOS* (**Figure 4h**), a major transcriptional regulator of T cell effector function(*52*). Notably, we also found increased accessibility at the *CHD5* locus (**Figure 4h**) and elevated *CHD5* mRNA expression (**Figure 4i**) upon CHD4 loss, suggesting that CHD4 supports the repression of the CHD5 locus. Together, these findings point to functional specificity among CHD family members within the NuRD complex.

To systematically connect changes in chromatin accessibility to transcriptional output, we constructed an enhancer-based gene regulatory network (eGRN) using the GRaNIE framework(*53*), integrating ATAC-seq and RNA-seq data to infer transcription factor (TF)-enhancer-gene relationships. The resulting network comprised 125 TFs that were connected to 8,723 regulatory regions and 9,748 target genes (**Figure 4j**). To quantify the extent to which TF–gene connections in the eGRN explain the differential gene expression between CHD4-KO and control T cells, we applied GRaNPA(*53*), which uses machine learning to estimate the contribution of individual TFs to the observed transcriptional changes. GRaNPA showed robust predictive performance in explaining differential gene expression (**Supplementary Figure 6i**). Examination of TF contributions identified NFYB, HINFP, ZBTB14, JUND, TAF1, JUN and SP4 as the regulators with the highest predictive importance on both day 4 and 6 post- activation (**Figure 4k**). Notably, this set included the AP-1 family TFs JUN, JUND, and FOSL1, which are central transcriptional drivers of effector T cell function(*52*, *54*). Consistent with their high importance scores, predicted target genes of these TFs exhibited coordinated expression changes between CHD4- KO and control T cells (**Figure 4l**). Specifically, predicted target genes of NFYB and HINFP showed increased expression upon CHD4 loss, whereas target genes of ZBTB14 and TAF1 were predominantly downregulated. GO term overrepresentation analysis of TF target genes revealed enrichment for biological processes relevant for T cell effector function (**Figure 4m, Supplementary Table 12**). As expected, targets of AP-1 family members were enriched for pathways related to translation and mitochondrial gene expression, and targets of master regulator TCF7 showed strong enrichment for lymphocyte differentiation programs. ZBTB14 targets were also significantly enriched for T cell differentiation-related processes (**Figure 4m**), suggesting a potential novel role for this factor in regulating differentiation-associated transcriptional programs downstream of CHD4.

Given that CHD4 loss alters transcriptional programs associated with T cell differentiation, we next asked whether the identified regulatory modules were also associated with differentiation states of the cancer-associated T cells. To this end, we applied the CHD4-KO eGRN framework to predict transcriptional changes between double-positive (tumor-CD39^POS^) and single-positive (tumor-CD39^NEG^) T cells derived from the NSCLC patient samples. Notably, several TFs that we identified as drivers in CHD4-KO T cells (**Figure 4k**) also strongly contributed to the transcriptional differences between tumor- CD39^POS^ and tumor-CD39^NEG^ cells. These included JUN, HINFP, ZBT14, TAF1, SP4, and NFYB (**Figure 4n**). Consistently, predicted target genes of these TFs showed prominent differential expression between the tumor-CD39^POS^ and tumor-CD39^NEG^ T cell populations (**Figure 4o**). Notably, the predicted target genes of TAF1 and ZBT14 were downregulated in CHD4-KO T cells (**Figure 4l**) but upregulated in tumor- CD39^POS^ relative to tumor-CD39^NEG^ T cells (**Figure 4o**), consistent with the increased CHD4 protein expression observed in tumor-CD39^POS^ T cells (**Supplementary Figure 4a**). A similar opposite pattern was observed for target genes of ETS1, which has previously been described as a gatekeeper for dysfunctional T cell differentiation(*13*). Together, these findings show that CHD4-dependent chromatin remodeling represses transcriptional programs linked to T cell effector differentiation. Importantly, these regulatory circuits are also operational in tumor-derived dysfunctional T cells.

### FASN deletion mitigates chronic TCR stimulation–induced T cell dysfunction

Having established a role for CHD4 in regulating effector-associated transcriptional programs, we next investigated the contribution of the metabolic enzyme FASN to CD8^+^ T cell function. To understand whether FASN influences the functional state of CD8^+^ T cells, we assessed surface protein and transcription factor expression using flow cytometry in FASN knock-out (FASN-KO) T cells compared to T cells transfected with a non-targeting (NT) Cas9 complex (**Figure 5a, Supplementary Figure 7a**). Aside from changes in CCR7 and CD8 expression, FASN-KO T cells exhibited minimal changes (**Figure 5b-c, Supplementary Figure 7b**), suggesting that FASN does not play a large role in T cell state under standard culture conditions.

**Figure 5.**
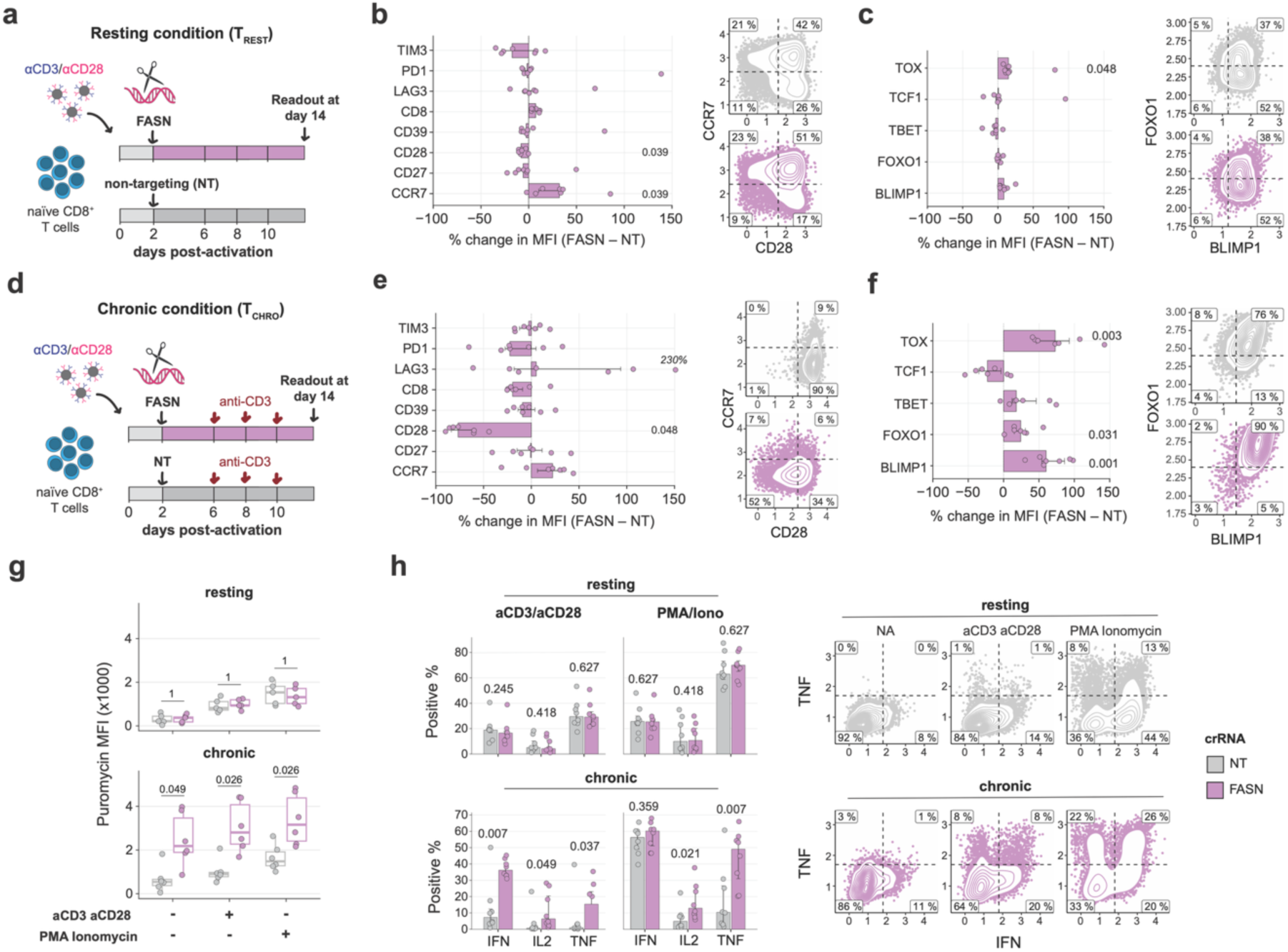
FASN deletion mitigates chronic TCR stimulation–induced T cell dysfunction. (**a-c**) Human naïve CD8^+^ T cells were activated with anti-CD3/CD28 dynabeads, nucleofected with *FASN* or non-targeting (NT) Cas9–guide RNA complexes at day 2 post-activation and analyzed at day 14 post-activation. (**a**) Experimental timeline. (**b-c**) Relative protein expression levels (FASN-deficient versus proficient) for indicated surface proteins (b) and transcription factors (c), measured by flow cytometry. Summarizing bar plots (left) and density plots of selected proteins (right) are shown. Dots indicate measurements from individual donors (n = 7), bars indicate group median, error bars indicate 25th/75th percentiles. (**d-f**) Activated CD8^+^ T cells nucleofected with *FASN* or non-targeting Cas9–guide RNA complexes were repetitively exposed to plate-bound anti-CD3 antibodies for 7 days (indicated with red arrows) and analyzed at day 14 post-activation. (**d**) Experimental timeline. (**e-f**) Relative protein expression levels (FASN-KO versus NT) for indicated surface proteins (e) and transcription factors (f), measured by flow cytometry. Summarizing bar plots (left) and density plots of selected proteins (right) are shown. Dots indicate samples obtained from individual donors (n = 7), bars indicate group median, error bars indicate 25th/75th percentiles. (**g**) Puromycin incorporation following 3-hour restimulation with anti-CD3/CD28 or PMA/ionomycin, with puromycin added during the last 10 minutes. Dots indicate measurements from individual donors (n = 6). Boxes show median and 25th/75th percentiles, whiskers extend to 1.5× the interquartile range. (**h**) Intracellular cytokine staining of indicated cytokines after a 3-hour re-stimulation with anti-CD3/CD28 antibodies or PMA/ionomycin in the presence of brefeldin A. Summarizing bar plots (left) and representative density plots (right) are shown. Dots indicate samples obtained from individual donors (n = 7), bars indicate group median, error bars indicate 25th/75th percentiles. Displayed graphs are representative of 2-4 independent experiments and contain aggregated data from 2 (**g**) or 3 (**b, c, e, f, h**) experiments. P values in all panels were determined by multiple paired two-sided Student’s t-tests with Benjamini–Hochberg FDR correction for multiple testing.

We therefore hypothesized that FASN may play a more prominent role during loss of function in CD8^+^ T cells. To test this, we established an in vitro model mimicking chronic TCR-stimulation (**Supplementary Figure 7c**). This model induces T cells (T_ChRO_) that phenotypically resemble dysfunctional T cells (e.g., TOX, PD-1, LAG3), display reduced cytokine production (TNF, IFNγ, IL-2), and exhibit diminished degranulation compared to ‘resting’ T cells (T_REST_). Notably, protein expression of both CHD4 and FASN was elevated in T_ChRO_ cells relative to T_REST_ cells, consistent with our proteomic findings in tumor-CD39^POS^ T cells (**Supplementary Figure 7d**). When we measured the effect of FASN deletion in this chronic TCR-stimulation model (**Figure 5d**), we observed more pronounced changes in surface protein and transcription factor expression in T_ChRO_ FASN-KO T cells compared to control T cells, suggesting that FASN is more important in T cells under stress conditions. Specifically, FASN-KO T_ChRO_ cells showed evidence of a more progenitor-like state, as indicated by increased expression of CCR7 and FOXO1 (**Figure 5e-f, Supplementary Figure 7e**). Paradoxically, the expression of TOX and BLIMP1 was also increased, while TCF1 and the co-stimulatory factor CD28 were reduced (**Figure 5e-f**), suggesting a partial uncoupling of dysfunction-related features in FASN-KO T cells. To assess whether these phenotypic changes resulted in alterations in cellular fitness, we performed a puromycin incorporation assay to measure global translation activity. Whereas the global protein synthesis rates were similar between FASN-KO and control populations in T_REST_ cells, T_ChRO_ FASN-KO cells exhibited substantially higher protein translation activity (**Figure 5g**). Increased protein synthesis capacity is required for high-level cytokine production(*55*). Consistent with the lack of changes in T cells under resting conditions (**Figure 5a-c**), FASN-KO T_REST_ cells displayed no significant differences in cytokine production in response to anti-CD3/CD28 or PMA/Ionomycin. However, FASN-KO T_ChRO_ cells maintained significantly higher production of TNF, IFNγ, and IL-2 compared to FASN-proficient T cells (**Figure 5h, Supplementary Figure 7f**), indicating that FASN deletion maintained the effector function of T cells specifically during chronic TCR-stimulation.

### FASN deletion protects against chronic TCR stimulation–induced mitochondrial dysfunction

We next sought to understand the molecular mechanisms underlying the protective effects of FASN deficiency. FASN is a key rate-limiting enzyme responsible for *de novo* lipogenesis, and lipid droplet accumulation has previously been associated with cancer-associated T cell dysfunction(*31*, *56*). To investigate whether the functional rescue observed in FASN-KO T cells was due to changes in lipid accumulation, we performed a BODIPY incorporation experiment to visualize the storage of cellular neutral lipids. Surprisingly, we did not observe consistent differences between FASN-KO and control T cells under resting or chronic T cell culture conditions (**Figure 6a**), indicating that FASN activity may not be the primary driver of bulk lipid accumulation in T cells.

**Figure 6.**
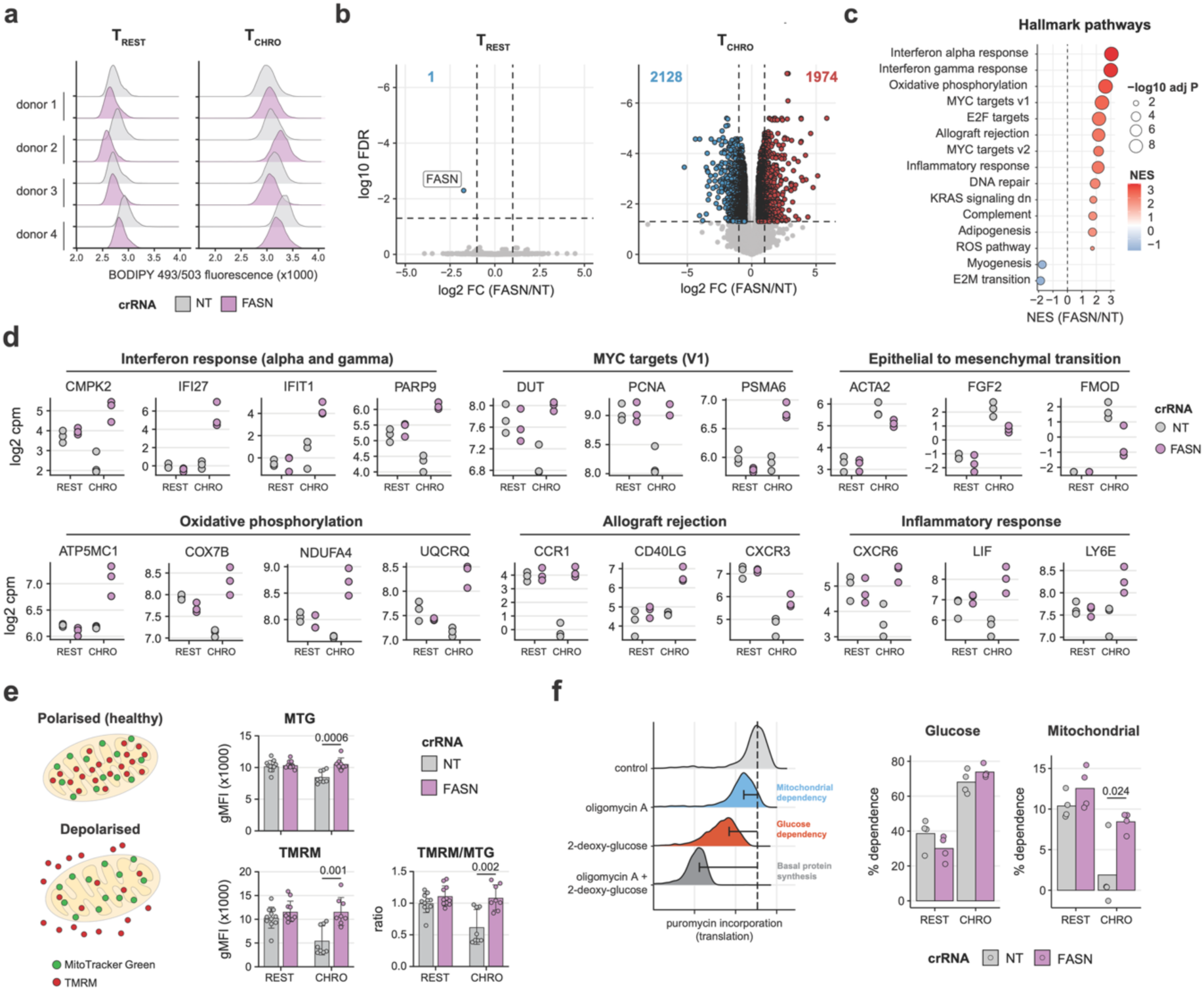
FASN deletion protects against chronic TCR stimulation–induced mitochondrial dysfunction. Activated resting (T_REST_) or chronically activated (T_CHRO_) FASN-KO or control (NT) CD8^+^ T cells (see Figure 5) were analyzed at day 14 post-activation. (**a**) BODIPY staining of indicated T cell populations measured by flow-cytometry. (**b-c**) RNA-sequencing analysis of FASN-KO and control T_REST_ and T_CHRO_ cells (n = 3) (**b**) Differential gene expression analysis. Transcripts with an FDR < 0.05 and fold change > 2 are highlighted. Bue and red numbers indicate the number of depleted and enriched transcripts, respectively. (**c**) Gene-set enrichment analysis (Hallmark pathways, Msigdb) of chronically stimulated FASN-KO versus control T cells. NES: Normalized enrichment score. (**d**) mRNA expression levels of representative genes from selected pathways shown in panel c. Dots indicate individual donors (n = 3). (**e**) MitoTracker Green (MTG) and tetramethylrhodamine methyl ester (TMRM) staining of indicated T cell populations, measured by flow-cytometry. Dots indicate individual donors (n = 8). (**f**) Dependence on glucose or mitochondrial metabolism assessed by the SCENITH assay. Dots indicate individual donors (n = 4). (**e, f**) Displayed graphs (**e-f**) are representative of 2-4 independent experiments (**a, e-f**), are aggregated from 3 experiments (**e**), or are derived from a single experiment (**b-d**). Barplots indicate group means, whiskers denote the standard deviation. P values were determined using edgeR’s quasi-likelihood F-test framework (**b**), permutation-based testing (**c**), or paired two-sided Student’s t-tests. Multiple-testing correction was performed using the Benjamini–Hochberg method (**b, c**) or the Bonferroni–Holm method.

To define the global molecular changes associated with FASN loss, we performed RNA sequencing on T_REST_ and T_ChRO_ cells in the presence or absence of FASN (**Supplementary Figure 7g-i**). Differential expression analysis confirmed that the effects of FASN-KO were largely restricted to T_ChRO_ cells as FASN was the only differentially expressed gene in T_REST_ cells (**Figure 6b, Supplementary Table 13**). Gene-set enrichment analysis indicated that FASN-KO T_ChRO_ cells displayed increased expression of genes involved in allograft rejection (e.g., CCR1, CD40LG, CXCR3), inflammatory responses (e.g., CXCR6, LIF, LY6E), and the interferon response (e.g., CMPK2, IFI27, IFIT1; **Figure 6c-d, Supplementary Table 14**). These findings are consistent with the enhanced translation capacity and functionality that we observed in FASN-KO T_ChRO_ cells (**Figure 5g-h**). Interestingly, genes involved in oxidative phosphorylation (e.g., ATP5MC1, COX7B, UQCRQ) were positively affected by FASN deletion in chronically stimulated T cells (**Figure 6c-d**). Because loss of mitochondrial fitness is a hallmark of T cell dysfunction(*57*), we next evaluated mitochondrial mass and membrane potential using MitoTracker Green (MTG) and TMRM staining (**Figure 6e**). Mirroring the mitochondrial deficiencies of cancer- associated dysfunctional T cells, T_ChRO_ cells exhibited both decreased MTG and TMRM signals compared to T_REST_ cells, together indicating reduced mitochondrial mass and membrane potential. Notably, and consistent with the observed transcriptional changes in FASN-KO T_ChRO_ cells, mitochondrial mass and function were both restored in these cells, indicating that loss of FASN counteracts the accumulation of mitochondrial defects that are associated with T cell dysfunction (**Figure 6e**).

Due to impaired mitochondrial respiration, dysfunctional T cells become increasingly dependent on aerobic glycolysis to meet their energetic demands(*57*, *58*). To functionally validate the MTG/TMRM findings and determine whether FASN deletion also restores metabolic flexibility in T_ChRO_ cells, we performed the SCENITH assay(*59*), which measures protein synthesis under metabolic inhibition as a proxy for cellular metabolic dependencies. SCENITH analysis demonstrated that FASN proficient T_ChRO_ cells predominantly utilized glycolysis and had a significantly reduced dependency on mitochondrial respiration (**Figure 6f, Supplementary Figure 7j**). In contrast, FASN-KO T_ChRO_ cells retained a level of dependency on oxidative phosphorylation that was comparable to that of T_REST_ (**Figure 6f, Supplementary Figure 7j**). Together, these findings demonstrate that the elevated expression of FASN during chronic TCR stimulation drives metabolic defects, and FASN deletion preserves mitochondrial fitness and limits the acquisition of transcriptional and phenotypic features that are associated with T cell dysfunction.

## Discussion

T cell dysfunction is a major contributor to cancer immunotherapy failure. However, our understanding of the cell intrinsic mechanisms that dampen T cell function in cancer remains limited. Here, we present a matched transcriptomic and proteomic map of cancer-associated dysfunctional and bystander T cells. In doing so, we identified a substantial discrepancy of mRNA- and protein-level fold-differences (Spearman r = 0.30–0.38) when comparing cancer-associated T cell populations. This discordance is consistent with previous studies of T cell populations isolated from peripheral blood(*34*, *60*). Such discrepancies may arise from two key factors: differences in the ability of the two methodologies to accurately quantify transcripts and proteins (technical causes)(*61–63*), or post- transcriptional events that independently regulate mRNA and protein abundance (biological causes)(*18*, *64*, *65*). Deconvolution of these two sources, for example by examining mRNA and protein sequence features (e.g., UTR motifs, codon usage, or stability determinants) associated with abundance discrepancies represents an important direction for future studies.

Effector T cell differentiation and acquisition of cytotoxic function are associated with dynamic rearrangements of epigenetic marks(*49–51*, *66*, *67*). For example, histones at effector-associated genes (e.g., *Ilr2a*, *Gzmb*, *Pdcd1*) are rapidly acetylated after T cell activation and subsequently de- acetylated during memory formation(*68*, *69*). We here identified the chromatin-remodeling factor CHD4 as a negative regulator of T cell function in NSCLC-derived dysfunctional T cells. CHD4 is a core component of the NuRD complex that comprises histone de-acetylases HDAC1/2. Previous studies have reported that restriction of HDAC activity increases genomic accessibility at effector-associated loci and thus enhances T cell function(*50*, *70*, *71*). Interestingly, we found that the protein abundance of the NuRD complex components remained comparable upon CHD4 deletion, likely attributed to the increased abundance of its neuronal-associated paralog CHD5 in CHD4-KO T cells. This increased CHD5 expression is, at least in part, epigenetically driven, given that the accessibility at its genomic locus is increased in CHD4-KO T cells. In line with these findings, CHD4 has been reported to repress lineage-determining genes(*36*, *72*). Moreover, CHD paralogs have previously been shown to repress one another’s expression(*73*). Despite this apparent compensation, genomic accessibility is strongly altered in CHD4-KO T cells, suggesting that CHD4 contributes to the target specificity of the NuRD complex. Target specificity is likely established through interactions with sequence-specific transcription factors, as CHD4 itself lacks sequence specificity(*74*, *75*). CHD4 is additionally part of the ChAHP complex, which modulates transcription by counteracting chromatin looping(*38*). Given the observed changes in chromatin accessibility upon CHD4 deletion, the ChAHP complex may also contribute to the observed phenotype in CHD4 KO T cells, in addition to the NuRD complex, although further studies are required to define its exact role in the effector T cell program.

We additionally identified elevated FASN expression in cancer-associated dysfunctional T cells. Increased fatty-acid synthase activity is a well-established feature of many cancers and promotes the accumulation of tumor-derived fatty acids that suppress infiltrating T cell function(*32*, *76*, *77*). In addition, the balance between fatty-acid oxidation, mitochondrial respiration, and glycolysis shapes T cell adaptation and effector function and is widely implicated in anti-cancer activity(*31*, *78*, *79*). In this study, we report that FASN acts as a T cell-intrinsic factor that limits T cell function, specifically in chronically activated T cells. Interestingly, FASN deletion did not alter neutral lipid content in T cells but unexpectedly preserved mitochondrial fitness of CD8^+^ T cells, which resulted in increased cytokine production capacity. Previous work has shown that FASN deficiency can increase cytosol-to-mitochondria citrate flux in tumor cells(*80*). A similar mechanism may operate in CD8^+^ T cells, potentially explaining the increased oxidative phosphorylation observed here through enhanced NADH availability for the mitochondrial electron transport chain. Together, these findings point to a potentially targetable axis in T cell dysfunction, as loss of oxidative phosphorylation and mitochondrial integrity are widely reported in cancer-associated T cells, and FASN inhibitors are currently being evaluated in clinical trials.

Collectively, our work demonstrates that integrating proteomic and transcriptomic analyses uncovers distinct molecular programs governing T cell function and differentiation that also shape dysfunctional states in tumor-infiltrating T cells.

## Methods

### Study Design

The objective of this study was to identify cell-intrinsic regulators of human cancer-associated T cell function by integrating proteomic and transcriptomic profiling with functional genetic perturbation. We performed proteomic and transcriptomic analyses of FACS-purified dysfunctional and bystander CD8⁺ tumor-infiltrating T cell populations isolated from treatment-naïve patients with non-small cell lung cancer, followed by CRISPR-Cas9-mediated loss-of-function studies, chromatin accessibility profiling, transcriptional analyses, and functional assays in primary human CD8⁺ T cells.

Sample sizes were based on the availability of patient material and on previous experience with the respective experimental approaches. Patient samples that met the predefined inclusion criteria were included in the study. Experimental samples were excluded only in cases of technical failure or when predefined quality control criteria for sequencing or mass spectrometry were not met. No statistical outliers were excluded unless stated otherwise.

Experiments were performed using biological replicates from independent patient samples or healthy donors, as indicated in the figure legends. Omics datasets were generated once per biological sample, whereas functional experiments were independently repeated with cells from multiple donors. Samples were not randomized, as experimental groups were defined by patient-derived T cell populations or by CRISPR-mediated gene perturbation. Investigators were not blinded during experiments or outcome assessment.

### Patient samples

The study was performed according to the Declaration of Helsinki (seventh revision, 2013), with consent of the Institutional Review Board of the Netherlands Cancer Institute/Antoni van Leeuwenhoek Hospital (NKI-AvL), Amsterdam, the Netherlands. Tumor and adjacent lung tissue of treatment-naïve, early-stage non-small-cell lung cancers patients was collected between April 2016 and November 2020(*25*, *26*). Briefly, within 4 hours after surgery, single cell suspensions were generated from isolated tumor or lung tissue by mechanical dissociation into small pieces, followed by enzymatic digestion in RPMI (Gibco) containing 30 IU/mL collagenase IV (Worthington), 12.5 μg/mL DNAse I (Roche), and 1% FBS (Bodego, Bodinco BV) for 45 minutes at 37 °C. Live cells were manually counted using trypan blue solution (Sigma) on a hemocytometer. Cells were cryo-preserved until further use.

### scRNAseq acquisition

Cryopreserved tumor digest was thawed and stained with anti-human CD3-APC (UCHT1, Biolegend), CD4-FITC (OKT4, Biolegend) and CD8-AF700 (SK1, Biolegend; **Supplementary Table 15**). Additionally, tumor digests were incubated with oligonucleotide barcode-conjugated antibodies (**Supplementary Table 15**) to detect antibody-derived tags (ADT) during single-cell RNA sequencing: anti-human CD4- TotalSeq-C0045 (SK3, Biolegend), CD8-TotalSeq-C0046 (SK1, Biolegend), CD25-TotalSeq-C0085 (BC96, Biolegend), CD279-TotalSeq-C0088 (EH12.2H7, Biolegend), CD39-TotalSeq-C0176 (A1, Biolegend), CD103-TotalSeq-C0145 (Ber-Act8, Biolegend) and CD127-TotalSeq-C0390 (A019D5, Biolegend). Cells were stained with propidium iodide (to identify live cells). Live single CD45^+^CD3^+^CD8^+^ T cells were sorted on a BD FACSAria™ Fusion Flow Cytometer (BD) and loaded into a Chromium single-cell sorting system (10x Genomics). Cells were kept on ice during all steps of the workflow. Gene expression and TCR library preparation was performed using the Chromium Next GEM Single Cell 5’ Kit v2, Chromium 5’ Feature Barcode Kit, and Chromium Single Cell Human TCR Amplification Kit (10x Genomics) according to the manufacturer’s instructions. Libraries were sequenced on Illumina NovaSeq 6000 sequencing systems (Illumina). Feature-barcode matrices were generated using the Cell Ranger software of the 10X Genomics Chromium™ pipeline.

### scRNAseq analysis

Initial scRNAseq data analysis was performed using the MetaCell(*28*) R package. Cells with less than 2,000 reads or a mitochondrial transcript fraction of lower than 0.075 were removed. Variable genes across the dataset were identified with a normalized variance/mean threshold at 0.12 and a down- sampled coverage threshold at 100, yielding 1,764 variable genes. These genes were subsequently used as anchors to calculate gene–gene correlations across the dataset. Genes with correlation coefficients above 0.1 were included. The obtained genes were then clustered into 20 separate gene-modules, and each gene-module was annotated manually for its biological function. Gene-modules falling in the categories ‘cell cycle’, ‘stress response’, ‘ribosomes’, and ‘histones’, were masked during MetaCell generation. Doublet detection was performed with the HTOdemux function of the Seurat package(*81*), setting the positive quantile at 0.99. For detailed settings for MetaCell generation, see https://github.com/kasbress/Proteomics_Cancer_Associated_Tcells. The MetaCell generation pipeline resulted in 85 MetaCells that were used for downstream analysis.

For correlation with previously published T cell dysfunction and cytotoxicity scores(*29*), centered log- ratio–normalized read counts were extracted for genes comprising each score. Scores were calculated per MetaCell (per patient) as the sum of read counts across the genes, normalized to the number of cells and scaled by 1,000. Antibody-derived tag (ADT) counts were normalized to counts per 10,000 across patients, and the median ADT signal per antigen was calculated for each MetaCell per patient. MetaCells comprising ≤5 cells per patient were excluded from the analysis. Spearman correlation coefficients were calculated per MetaCell per patient.

Pseudo-bulking of scRNA-seq counts was performed by grouping cells based on ADT signal for CD39 and CD103. Thresholds for defining CD103⁻CD39⁻ (DN), CD103⁺CD39⁻ (SP), and CD103⁺CD39⁺ (DP) populations were determined per patient (Supplementary Fig. 1c). TCR clonotypes were assigned at the single-cell level using the ‘filtered_contig_annotations.csv’ output from the 10x Genomics Cell Ranger pipeline and analyzed with the scRepertoire package. Unique CDR3 clonotypes were aggregated per population and patient, and repertoire diversity was quantified using the corrected Shannon index implemented in the iNEXT package(*82*).

### Cell sorting for LC-MS and bulk RNA sequencing

Cryopreserved tumor and adjacent tissue digests were thawed in a water bath at 37 °C. Cells were washed twice with pre-warmed (37 °C) RPMI containing 10% FBS (Bodinco) and 40 µg/mL DNAse I (Merck) and centrifugated at 250 × *g* for 8 minutes (acceleration 6, brake 6). Tissue digests were pre-incubated with Human TruStain FcX™ (Biolegend) and stained with anti-human CD3-FITC (SK7, Biolegend), CD8-AF700 (SK1, BioLegend), CD4-BV510 (SK3, Biolegend), CD25-AF647 (M-A251, Biolegend), CD27-BV605 (O323, Biolegend), CD69-BV421 (FN50, Biolegend), CD103-PE-Cy7 (Ber-ACT8, Biolegend), CD39-PE (A1, Biolegend) antibodies (**Supplementary Table 15)**, and Fixable Near-IR Dead Cell Stain (Invitrogen) in HBSS containing 1% FBS for 45 minutes at 4 °C. Samples were washed twice in HBSS by centrifugation at 250× *g* for 8 minutes (acceleration 6, brake 6) at 4 °C. Samples were sorted on a BD FACSAria™ III Cell Sorter. CD3^+^CD8^+^Near-IR^−^ T cells were divided into CD103^+^CD39^−^ and CD103^+^CD39^+^ T cell populations and sorted into Corning™ Costar™ Low Binding Plastic Microcentrifuge Tubes (FisherScientific), washed twice with ice-cold PBS, and snap-frozen in liquid nitrogen.

### Bulk RNAseq of cancer-associated T cells

Total RNA was isolated using the RNeasy Micro Kit (74004, Qiagen), including an on-column DNase digestion (79254, Qiagen), according to the manufacturer’s instructions. Quality and quantity of total RNA was assessed on the 2100 Bioanalyzer following manufacturer’s instructions “Agilent RNA 6000 Pico” (G2938-90046, Agilent Technologies). RNA samples were concentrated with Agencourt RNAClean XP beads (A63987), according to manufacturer’s instructions (Protocol 001298v001, Agencourt) with modifications. In short, 3 ng of total RNA per sample was brought to a volume of 30 μL by adding nuclease free water (AM9937, Ambion). 1.5 μL of Recombinant RNase inhibitor (2313A, Clontech) was added, and a 1.8x reaction volume RNAClean XP bead cleanup was performed with 18 μL RNAClean XP bead suspension and 36 μL RNAClean XP bead buffer (same solution without the beads) per sample. After 3 washes with 70% ethanol, bead pellets were dried at 37 °C for 3-5 minutes, until bead pellets start showing cracks. RNA was eluted by adding 4 μL of NF-H20 and vortexing. RNA library preparation was performed according to the published protocol “Full-length RNA-seq from single cells using Smart-seq2”(*83*) with modifications. In short, oligo dT primer hybridization was performed by adding Oligo dT mix (0.7 μL H_2_O, 0.1 μL RNAse inhibitor (40 U/μL), 0.1 μL dNTP mix (100 mM) and 0.1 μL Oligo-dT30VN primer (100 μM)) to the 4 μL sample-RNA bead suspension. Reverse transcription was performed as described, with the MgCl_2_ concentration adjusted to 10 mM. Template switching and 11 cycles pre-amplification of full length cDNAs with template switching oligos was performed using ISPCR primer at a final concentration of 0.08 μM. The amplified full-length cDNA was used for NGS library construction by Tagmentation for Illumina sequencing, using the Illumina Nextera XT DNA sample preparation kit (FC-131-1096, Illumina). RNA sequencing libraries were quantified and normalized based library QC data generated on the Bioanalyzer according to manufacturer’s protocols (G2938-90321, Agilent Technologies). A multiplex sequencing pool of all uniquely indexed RNA libraries was composed by equimolar pooling before paired end sequencing on the NovaSeq 6000 Illumina sequencing platform.

### TMT16 LC-MS of cancer-associated T cells

Cells were lysed in 50 μL lysis buffer (8 M urea, 100 mM Tris-HCl pH 8.0, 50 mM DTT) and sonicated (10 cycles of 30 seconds on/30 seconds off). Samples were incubated for 20 min at room temperature to ensure complete lysis. Alkylation was performed by adding iodoacetamide (IAA) to a final concentration of 50 mM and incubating for 15 minutes in the dark at room temperature. Proteins were enriched with SP3 beads(*84*). SP3 beads were prepared by combining Sera-Mag™ SpeedBeads (Sigma, GE65152105050250) and Sera-Mag™ Carboxylate-Modified Magnetic Particles (Sigma, GE45152105050250) at a 1:1 ratio. For each preparation, 20 μL of each bead type (total 40 μL) was mixed, washed three times with water, and resuspended in 100 μL water. Beads were stored at 4°C until use. To each sample, 5 μL of prewashed SP3 beads were added, followed by the addition of 1.4 volumes of acetonitrile and incubation for 20 minutes at room temperature. Samples were placed on a magnetic rack, and the supernatant was discarded. Beads were washed twice with 200 μL 70% ethanol and once with 200 μL acetonitrile, followed by brief air-drying.

Beads were resuspended in 50 μL 100 mM triethylammonium bicarbonate (TEAB) and digested with trypsin (1 μL, 0.1 μg/μL) for 2 hours at 37 °C with shaking (1500 rpm). Supernatant was collected, and beads were washed with an additional 50 μL TEAB, which was combined with the initial supernatant. Digestion was continued overnight at 37°C. Peptides were dried using a SpeedVac and resuspended in mass spectrometry-grade water to a final volume of 10 μL. TMT labeling was performed by adding 2.1 μL TMT reagent (0.5 mg dissolved in 24 μL acetonitrile) and incubated for 1 hour at room temperature. The reaction was quenched by adding hydroxylamine. Labeled samples were pooled and loaded onto C18 StageTips(*85*) for storage prior to further processing.

After elution from the stage tips, acetonitrile was removed with a SpeedVac. The remaining peptide solution was diluted with buffer A (0.1% FA) before loading. Peptides were separated on a 30 cm pico- tip column (75 µm ID, New Objective) in-house packed with 1.9 µm aquapur gold C-18 material (dr. Maisch) using a 240-minute gradient (5% to 80% Buffer B (80% ACN 0.1% FA)), delivered by an easy-nLC 1200 (Thermo), and electro-sprayed directly into a Orbitrap Eclipse Tribrid Mass Spectrometer (Thermo Scientific). The latter was set in data dependent mode with a cycle time of 3 seconds for both Faims CV settings (-45 V and -65 V), in which the full scan over the 380-1400 mass range was performed at a resolution of 120,000 and the automatic gain control target set to 400,000. The most intense ions (intensity threshold of 5000 ions, charge state 2-7) were isolated by the quadrupole with a 0.7 Da window and fragmented with a CID collision energy of 35%. The maximum injection time of the ion trap in turbo mode was set to 50 milliseconds with an AGC target of 10,000. Dynamic exclusion of 10 ppm was set on 90 seconds, including isotopes. TMT labels were analysed with an MS3 scan with synchronous precursor selection (SPS mass range of 400-1200) in the Orbitrap in enhanced resolution mode of 15,000, a range of 100-150 Th and a maximum injection time of 120 milliseconds. Precursors were isolated in the quadrupole with the same width (0,7 Da) as the MS2 but were fragmented with a normalized HCD collision energy of 55% and an AGC target of 100,000.

### LC-MS analysis of cancer-associated T cells

Raw mass spectrometry data were processed using Proteome Discoverer (Thermo Fisher Scientific) with the Sequest HT search engine. Spectra were searched against a human UniProt reference proteome database. Trypsin was specified as the protease, allowing up to two missed cleavages. Carbamidomethylation of cysteine residues and TMT labeling (TMTPro) at peptide N-termini and lysine residues were set as fixed modifications, while oxidation of methionine and protein N-terminal acetylation were included as variable modifications. Protein identification output from Proteome Discoverer was filtered to remove contaminants and retain only master proteins (designated by Proteome Discoverer). Proteins supported by at least two identified peptides were kept, and entries lacking quantified abundance values across all samples were excluded.

For principal component analysis (PCA), proteins were ranked based on variance across samples. The top 150 most variable proteins were selected for analysis. Abundance values were log2-transformed and arranged into a sample-by-protein matrix. PCA was performed using the ‘prcomp’ function implemented in R with scaling enabled. Only cancer-associated T cell populations were included in the analysis. The first two principal components were used for visualization.

Differential protein expression analysis was performed using the ‘limma’ R package. Linear models were fitted to log2-transformed protein abundances, with the different cancer-associated populations as factors. Log2 fold changes were estimated from model contrasts, and statistical significance was assessed using empirical Bayes moderation. Differentially expressed proteins (adjusted P < 0.05) were visualized using the ‘ComplexHeatmap’ R package. Protein abundances were arranged into a gene-by-sample matrix and z-score–scaled per protein. Values were capped at ±2 for visualization. Hierarchical clustering was performed using Euclidean distance for both rows and columns, and k-means clustering (k = 3) was applied to group proteins and samples. Gene set enrichment analysis (GSEA) was performed using the ‘clusterProfiler’ package in R. Genes were ranked by log2 fold change, as determined in the differential expression analysis (tumor-CD39^POS^ versus tumor-CD39^NEG^). Enrichment of Hallmark gene sets (MSigDB) was assessed using the GSEA function with default parameters. Adjusted P values were used to determine significance.

### Bulk RNAseq analysis of cancer-associated T cells

RNA-seq reads were aligned to the human reference genome (GRCh38/hg38) using the STAR aligner (v2.7.11b) with default parameters. The genome index was generated from the Ensembl GRCh38 primary assembly FASTA (release 113), matching the gene annotation. Aligned reads were output as coordinate-sorted BAM files. Gene-level read counts were quantified using featureCounts (Subread v2.0.2) based on the Ensembl GRCh38 (release 113) GTF annotation, with paired-end reads counted using default settings. PCA was performed on log2-transformed counts per million (CPM) values using the top 2,000 most variable genes (based on variance across samples). PCA was performed using the ‘prcomp’ function with scaling enabled. The first two principal components were used for visualization. Differential RNA expression analysis was performed using the ‘limma’ package, as described above for differential protein expression.

### scRNAseq re-analysis of the Salcher dataset

Single-cell RNA-sequencing datasets were obtained from the lung cancer atlas available through the CELLxGENE Discover platform(*86*). Cells annotated by the original study authors as “T cell regulatory”, “T cell CD4”, or “T cell CD8” were subsetted and reprocessed using the standard Seurat workflow, including normalization, identification of variable features, scaling, dimensionality reduction, neighborhood graph construction, and clustering. CD8⁺ T cells were subsequently isolated and annotated using ProjecTILs(*87*). Reference mapping was performed against the human CD8⁺ TIL reference atlas (CD8 TIL version 1)(*88*). Following ProjecTILs annotation, cells were reprocessed using the default Seurat pipeline prior to downstream analyses.

Next, studies included in the meta-analysis were filtered for those including material of at least 8 patients for which more than 200 single cells were sequenced. These studies were: ‘Chen_Zhang_2020’, ‘Guo_Zhang_2018’, ‘Kim_Lee_2020’, ‘Leader_Merad_2021_10x_3p_v2_beads’, ‘UKIM-V-2’. Gene expression data were pseudo-bulked per patient for each ProjecTILs-defined CD8⁺ T cell population using the ‘AggregateExpression’ function in Seurat. ProjecTILs populations were manually re-annotated based on marker expression. TCM/TEM populations were classified as tumor- CD39^NEG^ (ITGAE-high and ENTPD1-low), and TEX populations were classified as tumor-CD39^POS^ (ITGAE- high and ENTPD1-high). Differential expression analysis was performed using the ‘edgeR’ package. Lowly expressed genes were filtered using the ‘filterByExpr’ function (minimum count = 5; minimum proportion of samples = 0.5). Library sizes were normalized using the trimmed mean of M-values (TMM) method. A design matrix was specified to model differences between populations. Dispersion estimates were obtained using quasi-likelihood methods, and generalized linear models were fitted using ‘glmQLFit’. Differential expression between populations was assessed using quasi-likelihood F-tests (glmQLFTest).

### Generation of CRISPR Cas9–guide RNA complexes

Alt-R™ crRNAs targeting the genes of interest (**Supplementary Table 16**) and Alt-R™ ATTO550-labeled tracrRNA (Integrated DNA Technologies, IDT) were reconstituted to 100 µM in IDTE buffer (pH 7.5) according to the manufacturer’s instructions. As a control, Alt-R™ CRISPR-Cas9 Negative Control crRNA #1 (IDT) was used. crRNA and tracrRNA were mixed at equimolar ratios (32 pmol of each crRNA per reaction) and annealed by incubation at 95 °C for 5 minutes in a thermocycler, followed by gradual cooling to 20 °C at a ramp rate of 0.1 °C/s to allow duplex formation. Recombinant Alt-R™ S.p. Cas9 Nuclease V3 (IDT; 10 µg/µL) was added to the crRNA-tracrRNA duplex for at least 10 minutes at room temperature to form ribonucleoprotein (RNP) complexes.

### T cell culture

Peripheral blood mononuclear cells (PBMCs) from anonymized healthy donors were used in accordance with the Declaration of Helsinki (Seventh Revision, 2013) after written informed consent (Sanquin). PBMCs were isolated through Lymphoprep density gradient separation (Stemcell Technologies), followed by cryopreservation.

Naïve CD8^+^ T cells were isolated from PBMCs of individual donors using the BD IMag™ Human CD8 T Lymphocyte Enrichment Set (BD Biosciences), supplemented with 1 µg biotinylated anti-human CD45RO (UCHL1, BioLegend) per 1×10⁶ cells, following the manufacturer’s protocol, routinely achieving >90% purity. 100,000 naïve CD8^+^ T cells were activated for 3 days in tissue-culture treated flat-bottom 96-well plates using Dynabeads^™^ Human T-Activator CD3/CD28 for T Cell Expansion and Activation (Gibco). Cells were harvested and Dynabeads^™^ were removed using a BD IMag™ Cell Separation Magnet (BD Biosciences). T cells were cultured in T cell medium: RPMI (Gibco) supplemented with 10% FBS (Serana Europe), 100 U/mL penicillin, 100 µg/mL streptomycin, 25 mM HEPES (Gibco), non-essential amino acids (Gibco), 10 mM sodium pyruvate (Gibco), 50 IU/mL rhIL2 (Protech), 5 ng/mL rhIL15 (Gibco) and 10 ng/mL IL7 (Gibco) at 37°C and 5% CO_₂_. Fresh T cell medium was added to T cell cultures every 2–3 days, and cells were kept at a density of 0.5–2.0 × 10^⁶^ cells/mL. For nucleofection with CRISPR–Cas9 ribonucleoprotein (RNP) complexes, T cells were activated for 2 days. 100,000–500,000 activated T cells were harvested per condition and washed twice with PBS by centrifugation at 300 × *g* for 5 minutes. Cells were resuspended in 20 µL TheraPEAK® P3 Primary Cell Nucleofector® Solution (Lonza) and mixed with 16–25 pmol CRISPR–Cas9 RNP and 32 pmol Alt-R™ Cas9 Electroporation Enhancer (IDT). Cell suspensions were transferred to a 16-well Nucleocuvette® Strip (Lonza) and nucleofected using a 4D-Nucleofector® X Unit (Lonza) with pulse code EH-115. Pre-warmed (37 °C) T cell medium was added immediately after nucleofection. Cells were cultured for another 24 hours before Dynabeads™ were removed and T cells cultured as described above. Fresh T cell medium was added to T cell cultures every 2–3 days, and cells were kept at a density of 0.5-2.0×10⁶ cells/mL.

### Western blotting

4 days after electroporation, CD8⁺ T cells were harvested and 0.4 × 10⁶ cells were snap-frozen per condition and stored at −80 °C for up to two weeks. Cell pellets were lysed for 10 min on ice in 40 µL RIPA buffer (Thermo Fisher Scientific) supplemented with 1% (v/v) Halt™ Protease Inhibitor Cocktail (Thermo Fisher Scientific) and 500 mM dithiothreitol. The soluble fraction was separated and cleared by centrifugation at 13,000g for 10 minutes. For histone and HDAC immunoblotting, RIPA buffer was supplemented with 1% (v/v) Halt™ Protease Inhibitor Cocktail, 0.1666 U/µL Benzonase® Nuclease (Sigma Aldrich), and 10 mM MgCl₂ and 10 mM CaCl₂ to facilitate chromatin digestion. Samples were mixed with 5% (v/v) LDS sample buffer (Invitrogen) and denatured for 5 minutes at 95 °C. Proteins were separated on 4–12% Novex Bis-Tris mini gels (Invitrogen) using MOPS running buffer (50 mM MOPS, 50 mM Tris-HCl pH 7.7, 0.1% SDS, 1 mM EDTA) and transferred onto nitrocellulose membranes using the iBlot 2 Dry Blotting System (Invitrogen). Membranes were blocked for 1 hour at room temperature in TBST containing 2.5% BSA. Primary antibodies (**Supplementary Table 15**) were added (1:1000) to TBST containing 2.5% BSA and incubated overnight at 4 °C. Blots were washed 5x for 5 minutes and membranes were incubated for 1 hour at room temperature with secondary α-mouse or α-rabbit HRP antibodies at 0.1% (v/v) in TBST containing 2.5% BSA. Blots were washed 5x for 5 minutres in TBST and proteins were detected using the Thermo Scientific™ Pierce™ ECL Western Blotting Substrate (Thermo Fisher Scientific). Densitometry analyses were performed using Fiji v1.8.0.

### Re-stimulation analysis

For restimulation experiments, 100,000 T cells per condition were reactivated in flat-bottom 96-well plates at the indicated time points using anti-CD3 PeliCluster (Sanquin Reagents) and anti-CD28 (CD28.2; eBioscience) stimulation at 1 µg/mL each, or 10 ng/mL phorbol 12-myristate 13-acetate (PMA) and 1 µM ionomycin. Cells were stimulated for 3 hours at 37 °C and 5% CO₂. For staining of CD107a, anti-human CD107a-BUV395 (H4A3; BD Biosciences) was added immediately during activation. Brefeldin A Solution 1000x (eBioscience) was added after the first 15 minutes of the stimulation period to allow intracellular accumulation of cytokines. Following restimulation, cells were processed for downstream assays, including intracellular cytokine staining and flow cytometric analysis as described above.

For global translation assays, 5 µg/mL puromycin was added during the final 10 minutes of culture to allow incorporation into nascent peptides as a measure of protein synthesis. Cells were then processed as described above for intracellular staining of cytokines, using an anti-puromycin-AF647 (12D10; Merck) antibody for detection by flow cytometry.

### Co-culture assay

Mel888 cells expressing a membrane-thethered single-domain variable fragment OKT3 antibody(*48*) were cultured in IMDM (Gibco) supplemented with 8% FBS (Bodinco), 100 U/mL penicillin and 100 µg/mL streptomycin at 37 °C and 5% CO₂. Cells were passaged every 2-3 days at a 1:10 split ratio. For functional co-culture assays, 15,000 Mel888 target cells were seeded per well in flat-bottom 96- well plates in IMDM supplemented with 8% FBS, 100 U/mL penicillin and 100 µg/mL streptomycin. Cells were centrifuged at 10 × *g* for 1 minute without brake, and incubated for 4-5 hours at 37 °C and 5% CO_2_. 150 µL of medium was removed, and 15,000 CD8⁺T cells in 150 µL T cell medium (described above) containing 50 U/mL IL-2 were added per well. Plates were centrifuged at 10 × *g* for 1 minute without brake and incubated overnight at 37 °C and 5% CO_2_. For cytokine analysis, Brefeldin A Solution 1000x (eBioscience) was added to the media 20 minutes after the start of coculture to allow accumulation of cytokines for downstream analysis.

### Flow cytometric analysis

100,000 T cells were harvested from cultures at indicated time points and washed with PBS. For CCR7 detection, cells were incubated with anti-CCR7-AF647 (3D12; BD Biosciences) in T cell media for 30 minutes at 37 °C and 5% CO₂. Subsequently, surface staining was performed for 15 minutes at room temperature in PBS containing 1% FBS (Bodinco) and LIVE/DEAD™ Fixable Near-IR (Invitrogen), to exclude dead cells, and CD3-FITC (SK7; BioLegend), CD4-BV510 (SK3; BioLegend), CD8-AF700 (SK1; BioLegend), CD25-AF647 (M-A251; BioLegend), CD27-BV605 (O323; BioLegend), CD28-PE-Cy7 (CD28.2; BioLegend), CD39-PE (A1; BioLegend) or CD39-BUV395 (A1; BD Biosciences), CD69-BV421 (FN50; BioLegend), CD103-PE-Cy7 (Ber-ACT8; BioLegend), PD-1-BV421 (EH12.1; BD Biosciences), TIM- 3-BB515 (7D3; BD Biosciences), LAG-3-PE (11C3C65; BioLegend), and CD137-BV605 (4B4-1; BioLegend) antibodies (**Supplementary Table 15**), as indicated.

Intracellular staining with antibodies against IFNγ-PE (B27; BioLegend), IL-2-APC (MQ1-17H12; BioLegend), TNF-PE-Cy7 (MAb11; BD Biosciences), granzyme B-AF700 (GB11; BD Biosciences), and was performed after fixation and permeabilization using the BD Cytofix/Cytoperm™ Fixation/Permeabilization Kit (BD Biosciences) according to the manufacturer’s instructions. For intra- nuclear staining with FOXO1-AF488 (C29H4; Cell Signaling Technology), T-bet-BV421 (4B10; BioLegend), TCF1-PE (C63D9; Cell Signaling Technology) and TOX-APC (REA473; Miltenyi Biotec) antibodies, cells were processed using the Foxp3/Transcription Factor Staining Buffer Set (eBioscience) following the manufacturer’s protocol. After washing, samples were resuspended in PBS containing 1% FBS and acquired on a BD LSRFortessa (BD Biosciences) flow cytometer. Data were analyzed using FlowJo software v10.10.0 (TreeStar).

### Cell cycle and Apoptosis analysis

Cell proliferation was measured by intracellular staining of Ki67-AF647 (B56; BD Biosciences). Cells were harvested, washed with PBS, and fixed using BD Cytofix/Cytoperm Kit (BD Biosciences) as described above. After washing, cells were resuspended in PBS containing 1% FBS (Bodinco) containing propidium iodide 1:20 (PI; Invitrogen) and incubated for 15 minutes prior to acquisition.

Cell viability and apoptosis was assessed by staining T cells with Annexin V-APC (1:50, BD Pharmingen) in Annexin V Binding Buffer (BioLegend)™) for 15 minutes at room temperature. Cells were washed twice and resuspended in Annexin V binding buffer. Immediately prior to flow cytometric acquisition, PI was added at 1:20 dilution in binding buffer.

### Pulldown experiments HDAC1 and CHD4

Co-immunoprecipitation experiments were performed using the Pierce Classic Magnetic IP/Co-IP Kit (Thermo Fisher Scientific) according to the manufacturer’s instructions. Briefly, 1.5 × 10⁶ CD8⁺ T cells per condition were lysed in IP Lysis/Wash Buffer and clarified by centrifugation. For each immunoprecipitation, 0.4 µg of polyclonal antibody against CHD4 (ProteinTech), HDAC1 (ProteinTech), or Rabbit IgG control polyclonal antibody (ProteinTech) was added to the lysate and incubated for 1 hour at room temperature, followed by overnight incubation at 5 °C to allow immune complex formation. Subsequent binding to Protein A/G magnetic beads was performed according to the manufacturer’s protocol. Washing conditions were adjusted to prepare samples for LC-MS analysis: After overnight incubation, beads were washed five times using a wash buffer consisting of 0.025 M Tris, 0.15 M NaCl, and 0.001 M EDTA.

Beads were resuspended in 50 μL 1M Urea/100 mM Tris-HCl pH 8 (Invitrogen). Proteins were reduced using 10 mM dithiothreitol (Thermofisher Scientific) for 20 minutes at 25°C, and reduced cysteine residues were alkylated with 50 mM iodoacetamide (Thermofisher Scientific) for 10 minutes in the dark at 25°C. Proteins were digested on beads using 250 ng MS-grade Trypsin/LysC (Thermofisher Scientific) for 2 hours at 25°C. The supernatant fraction was collected after digestion, whereas beads were washed with 50 μl 1M Urea/10 mM DTT in 100 mM Tris-HCl (pH=7.5) for 5 minutes at 25°C. Supernatants from both fractions were combined, and proteins were further digested overnight at 25°C using 100 ng Trypsin/LysC. Tryptic digests were acidified by adding 11 μl 10% trifluoroacetic acid (Thermofisher Scientific), loaded on Evotips (Evosep) according to manufactur’s guidelines and separated on an 8 cm × 150 μm, 1.5 μm Performance Column (EV1115 from EvoSep) with an Evosep One liquid chromatography (LC) system (Evosep) using the 60 samples per day gradient. Peptides were ionized and electrosprayed into a TimsTOF HT mass spectrometer (Bruker). Data was acquired in DIA-PASEF mode, using an MS1 scan range of 100-1700 m/z. For MS2 acquisition 32 pyDIAID system- optimized23 windows were used, with a cycle time of 1.80 seconds, and a mass and ion mobility range from 400.2-1500.8 m/z and 0.70-1.50 1/k0 respectively. Collision energy used was 20.00 eV at 0.6 1/k0 and 59 eV at 1.60 1/k0. Raw mass spectrometry data files were processed using DIA-NN (version 2.3.1). Proteins and peptides were detected by querying the human reference proteome (UP000005640, release 2025.03.06). Standard settings were used, using a generated library-based spectra search. Maximum number of variable modifications was set at 1, and peptide length range to 6-30 amino acids. Data was analyzed using R 4.4.3 / Rstudio (2026.04.0). Detected proteins were filtered for proteotypic peptides and at least one unique peptide per protein. LFQ values were transformed to a log2 scale. Missing values were imputed by normal distribution (width = 0.3, shift = 1.8), assuming these proteins were close to the detection limit. Label free statistical analyses were performed using LIMMA, a Benjamin Hochberg-adjusted p < 0.05 and |log2 fold change| > 1 was used as significance threshold.

### RNA sequencing analysis of CHD4- and FASN-KO T cells

RNA-Seq was performed by Plasmidsaurus using Illumina Sequencing technology with custom analysis and annotation. RNA-seq FASTQ files were first assessed for quality using FastQC (v0.12.1). Adapter removal and quality filtering were performed using fastp (v0.24.0), including poly-X tail trimming, 3ʹ end quality trimming, a minimum Phred quality score threshold of 15, and a minimum read length cutoff of 50 bp. Filtered reads were aligned to the human reference genome (GRCh38/hg38, release 115) using the STAR aligner (v2.7.11) with default parameters, including removal of non-canonical splice junctions and reporting of unmapped reads. Aligned reads were coordinate-sorted using samtools (v1.22.1). PCR and optical duplicates were removed using UMI-based deduplication with UMIcollapse (v1.1.0).

Alignment quality metrics, including strand specificity and genomic feature distribution, were evaluated using RSeQC (v5.0.4) and Qualimap (v2.3). Quality control metrics were aggregated into a comprehensive report using MultiQC (v1.32). Gene-level expression quantification was performed using featureCounts (Subread v2.1.1) with strand-specific counting, fractional assignment of multi- mapping reads, and annotation based on exons and 3ʹ UTR features, summarized at the gene level (gene_id) using the Ensembl GRCh38 (release 115) GTF annotation. Gene counts were further annotated with gene biotype and additional metadata extracted from the reference annotation.

For downstream analyses, counts were normalized using trimmed mean of M-values (TMM), and sample–sample correlations were computed using Pearson correlation. Differential gene expression analysis was performed using edgeR (v4.0.16), including filtering of lowly expressed genes using ‘edgeR::filterByExpr’ with default settings. PCA was performed on log2-transformed counts per million (CPM) values using all or the top 2,000 most variable genes. Gene set enrichment analysis (GSEA) was performed using the ‘clusterProfiler’ package in R. Genes were ranked by log2 fold change, as determined in the differential expression analysis (tumor-CD39^POS^ versus tumor-CD39^NEG^), and enrichment of Hallmark gene sets (MSigDB) was assessed using the GSEA function with default parameters. Adjusted P values were used to determine significance.

### Assay for Transposase-Accessible Chromatin using sequencing (ATAC-Seq)

Chromatin accessibility profiling was performed using a modified version of the protocol from(*89*), using in-house produced Tn5 from the EMBL Protein Expression and Purification facility(*90*). Tn5 was pre-loaded with Nextera-compatible sequencing adapters as previously described. Briefly, 80,000 CD8⁺ T cells were harvested for each condition, and nuclei were isolated by incubating on ice for 4 minutes in lysis buffer: 0.1% NP-40 (Thermo Scientific), and 0.01% digitonin (Invitrogen). Nuclei were counted after lysis, and tagmentation was carried out using 20,000 nuclei per sample, incubating at 37 °C for 30 minutes shaking at 500 rpm, using assembled Tn5 in tagmentation buffer: 18.8% DMF (Sigma-Aldrich), 11.8 mM Mg-acetate, 77.6 mM K-acetate, 38.8 mM Tris-acetate, 0.12% NP-40, and EDTA-free Halt™ Protease Inhibitor Cocktail (Thermo Scientific). Immediately following transposition, DNA was purified using the MinElute PCR Purification Kit (Qiagen), and libraries were PCR-amplified using NEBNext High-Fidelity 2X PCR Master Mix using combinatorial dual index primers (P5 and P7). Final libraries were size selected sequentially using SPRIselect beads (Beckman Coulter), applying a 1:1.2 and 1:0.6 sample to beads ratio. Quality and fragment distribution were assessed on the 2100 Bioanalyzer system using the High Sensitivity DNA Kit (Agilent). Final libraries were sequenced on a Novaseq X (10B, single lane, 300 cycles) at the Utrecht Sequencing Facility (USEQ, Utrecht, The Netherlands).

### ATAC-Seq analysis

Raw ATAC-sequencing data was processed with an ATAC-Seq data processing Snakemake pipeline as previously described(*91*). Raw reads were first assessed with FastQC (http://www.bioinformatics.babraham.ac.uk/projects/fastqc/), trimmed using Trimmomatic(*92*) (MINLEN cutoff 20), and aligned to the GRCh38/hg38 reference genome using Bowtie2(*93*) (“--very- sensitive”, maximum fragment length 2000 bp). Properly paired reads with MAPQ ≥10 were retained using SAMtools(*94*), and PCR duplicates were identified and removed using Picard MarkDuplicates (http://broadinstitute.github.io/picard). Reads containing insertions, deletions or soft-clipping were excluded based on CIGAR strings. Peaks were called using MACS2(*95*) with the parameters “—nomodel --shift -100 --extsize 200 -q 0.01”, and read counts per peak were quantified using featureCounts(*96*). Downstream analysis was performed in R using DiffBind(*97*) and DESeq2(*98*). A consensus peak set was generated from peaks detected in at least three samples, and reads were counted over summit-centered 250 bp regions. Differential accessibility analysis was performed with DESeq2 using a design controlling for donor effects (∼ donor + condition_timepoint). Differentially accessible regions were identified for CHD4-deficient versus control CD8⁺ T cells separately at day 4 and day 6. Peaks with adjusted P < 0.05 and absolute log2 fold change > 0.58 were considered differentially accessible. Peaks were annotated to genomic features and nearby genes using ChIPseeker(*99*) with TxDb.Hsapiens.UCSC.hg38.knownGene and org.Hs.eg.db databases, using a TSS window of ±3 kb.

### Gene regulatory network analysis

Gene regulatory network inference was performed using GRaNIE(*53*) Raw ATAC-seq counts from the consensus peak set and raw RNA-seq counts (obtained as described above) were supplied to GRaNIE using the hg38 genome assembly. ATAC-seq counts were normalized using DESeq2 size factors, and RNA-seq counts were quantile-normalized using limma. Transcription factor binding sites were overlapped with accessible regions using motifs from the HOCOMOCOv12 database(*100*). Peaks were filtered to retain regions with normalized mean accessibility ≥5, located on retained chromosomes. TF-peak and peak-gene connections were inferred using Pearson correlation, with peak-gene links restricted to genes within 250 kb of each peak. The final gene regulatory network was filtered using FDR < 0.2 for both TF-peak and peak-gene connections and retained protein-coding and lincRNA genes. GRaNPA was used to prioritize transcription factors associated with CHD4-dependent transcriptional changes. Filtered GRaNIE TF-gene connections were converted into a TF-target gene matrix, and DESeq2-derived RNA-seq log2 fold changes (obtained as described above) for CHD4-deficient versus control cells at day 4 and day 6 were used as gene-level input metrics. Only genes with adjusted P < 0.05 were included in the GRaNPA analysis. Gene Ontology (GO) enrichment analysis of transcription factor regulons was performed using the R package clusterProfiler(*101*). Gene sets of interest were tested for enrichment of Biological Process terms using the enrichGO function with the org.Hs.eg.db annotation database. Enrichment was assessed using a hypergeometric test, with all genes included in the analysis used as background. P-values were adjusted for multiple testing using the Benjamini– Hochberg method, and GO terms with adjusted P < 0.05 were considered significant. Redundant GO terms were reduced using the simplify function (cutoff = 0.5, by = "p.adjust", select_fun = min) in clusterProfiler.

### Chronic model

To model chronic TCR stimulation-induced T cell dysfunction, naïve CD8⁺ T cells were activated and cultured as described above. From day 6 post-activation onward, T cells were subjected to repeated TCR stimulation. Non-tissue-culture treated flat-bottom 24-well plates were coated with 2 µg/mL Ultra-LEAF™ anti-human CD3 antibody (HIT3a, BioLegend) for 3 hours at 37°C and used for T cell stimulation. Cells were restimulated with a freshly anti-CD3-coated plate every 2–3 days for a total of 7 days. Flow cytometry assays were performed as described above. For the assays described below, 50,000–100,000 FASN-proficient/deficient T_REST_ and T_ChRO_ cells (day 12-14 post-activation) were seeded 96-well v-bottom plates in RPMI (Gibco) supplemented with 10% FBS (Serana Europe), 100 U/mL penicillin, 100 µg/mL streptomycin, 25 mM HEPES (Gibco), 50 IU/mL rhIL2 (Protech), 5 ng/mL rhIL15 (Gibco) and 10 ng/mL IL7 (Gibco). Cells were incubated for at least 2 hours at 37°C and 5% CO₂ prior to the assay.

### Mitotrackers

50,000–100,000 T_REST_ and T_ChRO_ cells (FASN-KO and NT) were seeded on tissue-culture treated 96-well v-bottom plates at least 2 hours before the assay. As a positive control for the TMRM stain, mitochondria were hyperpolarized by pre-treating cells with 1 µM oligomycin A (MedChemExpress) for 10 minutes at 37 °C. As a negative control, mitochondria were de-polarized by pre-treating samples with 50 µM Carbonyl cyanide 3-chlorophenylhydrazone (CCCP; ThermoFisher). Samples were incubated with 40 nM MitoTracker™ Green FM (ThermoFisher) and 5 nM Image-iT™ TMRM Reagent (ThermoFisher) for 30 minutes at 37°C, washed with ice-cold PBS containing 1% FBS and stained on ice for 5 minutes with anti-human CD8-BUV805 (SK1, BioLegend) and Fixable Near-IR Dead Cell Stain (Invitrogen). Next, samples were washed twice with PBS containing 1% FBS and measured on a BD LSRFortessa™ Cell Analyzer (BD).

### BODIPY

50,000–100,000 T_REST_ and T_ChRO_ cells (FASN-KO and NT) were seeded on tissue-culture treated 96-well v-bottom plates at least 2 hours before the assay. BODIPY™ 493/503 (ThermoFisher) was pre-diluted to 10 µM in T cell media (without FCS) and mixed vigorously on a vortex to emulsify the solution. This emulsion was immediately added to the T cells to a final concentration of 5 µM and incubated at 37 °C for 30 minutes. Cells were washed with ice-cold PBS containing 1% FBS and stained for 5 minutes with anti-human CD8-AF700 (SK1, BioLegend) and Fixable Near-IR Dead Cell Stain (Invitrogen). Next, samples were washed twice with PBS containing 1% FBS and measured on a BD LSRFortessa™ Cell Analyzer (BD).

### SCENITH

Single cell energetic metabolism by profiling translation inhibition (SCENITH) was performed as described(*59*). In short, 50,000–100,000 FASN-proficient/deficient T_REST_ and T_ChRO_ cells (day 14 post- activation) were seeded on tissue-culture treated 96-well v-bottom plates in RPMI (Gibco) supplemented with 10% FBS (Serana Europe), 100 U/mL penicillin, 100 µg/mL streptomycin, 25 mM HEPES (Gibco), 50 IU/mL rhIL2 (Protech), 5 ng/mL rhIL15 (Gibco) and 10 ng/mL IL7 (Gibco). Cells were incubated for at least 2 hours at 37 °C and 5% CO₂ prior to the assay. Cultures were treated for 15 minutes with either 100 mM 2-deoxy-D-glucose (MedChemExpress), 1 µM oligomycin A (MedChemExpress), both inhibitors combined, or 2 µg/mL harringtonine. A DMSO-treated sample (matched v/v) was included as a control. Puromycin (10 µg/mL) was then added to all samples for an additional 25 minutes. Cells were washed in ice-cold PBS containing 1% FBS and stained for 5 minutes with anti-human CD8-AF700 (SK1, BioLegend) and Fixable Near-IR Dead Cell Stain (Invitrogen). Cells were fixed using the eBioscience™ Foxp3/ Transcription Factor Staining Buffer Set (Thermo Fisher) according to the manufacturer’s protocol, followed by blocking for 10 minutes at room temperature in permeabilization buffer containing 20% FBS. Samples were incubated for 1 hour at 4 °C with anti- puromycin antibody (12D10, Merck), washed twice with permeabilization buffer, and measured on a BD LSRFortessa™ Cell Analyzer (BD).

### Statistical analyses

All statistical analyses were performed either in R using the rstatix package or in python using the scipy package. Statistical tests, sample sizes, and definitions of biological and technical replicates are specified in the corresponding figure legends. All statistical tests were two-sided, and P < 0.05 was considered statistically significant. Where appropriate, P values were adjusted for multiple testing using the Benjamini–Hochberg or Holm correction. Differential protein expression was assessed using limma with empirical Bayes moderation, differential gene expression using edgeR’s quasi-likelihood F-test framework, and gene set enrichment analysis using clusterProfiler with permutation-based testing. Omics data were processed and normalized using standard methods as described in the corresponding Methods sections. Data are presented as mean ± SD or as boxplots indicating the median, interquartile range, and 1.5 × interquartile range, as specified in the figure legends.

## Supplementary Materials

Supplementary Figure 1–7

Supplementary Table 1–16

Supplementary Data 1

## Acknowledgements

We acknowledge the Utrecht Sequencing Facility (USEQ) for providing sequencing service and data (ATAC-seq). USEQ is subsidized by the University Medical Center Utrecht and The Netherlands X- omics Initiative (NWO project 184.034.019). We thank Suzanne Castenmiller for assistance with sample inclusion of cryopreserved material. We thank the Simon Tol and Erik Mul staff for technical support during fluorescence-activated cell sorting.

## Funding

European Research Council Consolidator grant Printers 817355 (MCW)

The Landsteiner Foundation for Blood Transfusion Research LSBR 2202 (MCW)

The MSCA-ITN grant RBP-REGUNET 101073094 (MCW)

Oncode Institute (MCW)

The Cancer Center Amsterdam research grant CCA2023-9-91 (KB)

The LSBR Early Career Grant LSBR ECG-2404 (NHV)

## Author contributions

Conceptualization: KB, ŽH, NHS, SS, MV, MCW

Methodology: KB, ŽH, NHS, SS, CGS, AJH, CZ, ŽM, RV, MN, JS, WS, MCW

Investigation: KB, ŽH, NHS, SS, CGS, MNM, AG, SK, AJH, CZ, ŽM, RV, MN, JS

Formal analysis: KB, ŽH, NHS, SS, MNM, NK, SK, AJH, CZ

Visualization: KB, ŽH, NHS, MNM, AJH

Data curation: KB, ŽH, NHS, SS

Resources: RvE, KM, KH, WSMET

Funding acquisition: KB, MV, MCW

Supervision: WS, MV, MCW

Writing – original draft: KB, ŽH, NHS

Writing – review & editing: KB, ŽH, SS, MV, MCW

## Data availability

The mass spectrometry proteomics data have been deposited to the ProteomeXchange Consortium via the PRIDE partner repository^29^ with the dataset identifier PXD079142 (Reviewer token: **EjY0lWfnyGuy**). The RNA sequencing and ATAC sequencing data from healthy donor material have been deposited on the European Nucleotide Archive (ENA) with the dataset identifier PRJEB114673. RNA sequencing data from NSCLC patient material has been deposited on the European Genome-phenome Archive (EGA) with the dataset identifier EGAS50000001886. All data needed for the reproduction of the figures presented in the work have been deposited to Zenodo (doi: 10.5281/zenodo.20713893). All analyses pipelines have been deposited at https://github.com/kasbress/Proteomics_Cancer_Associated_Tcells.

---

**Supplementary Figure 1.**
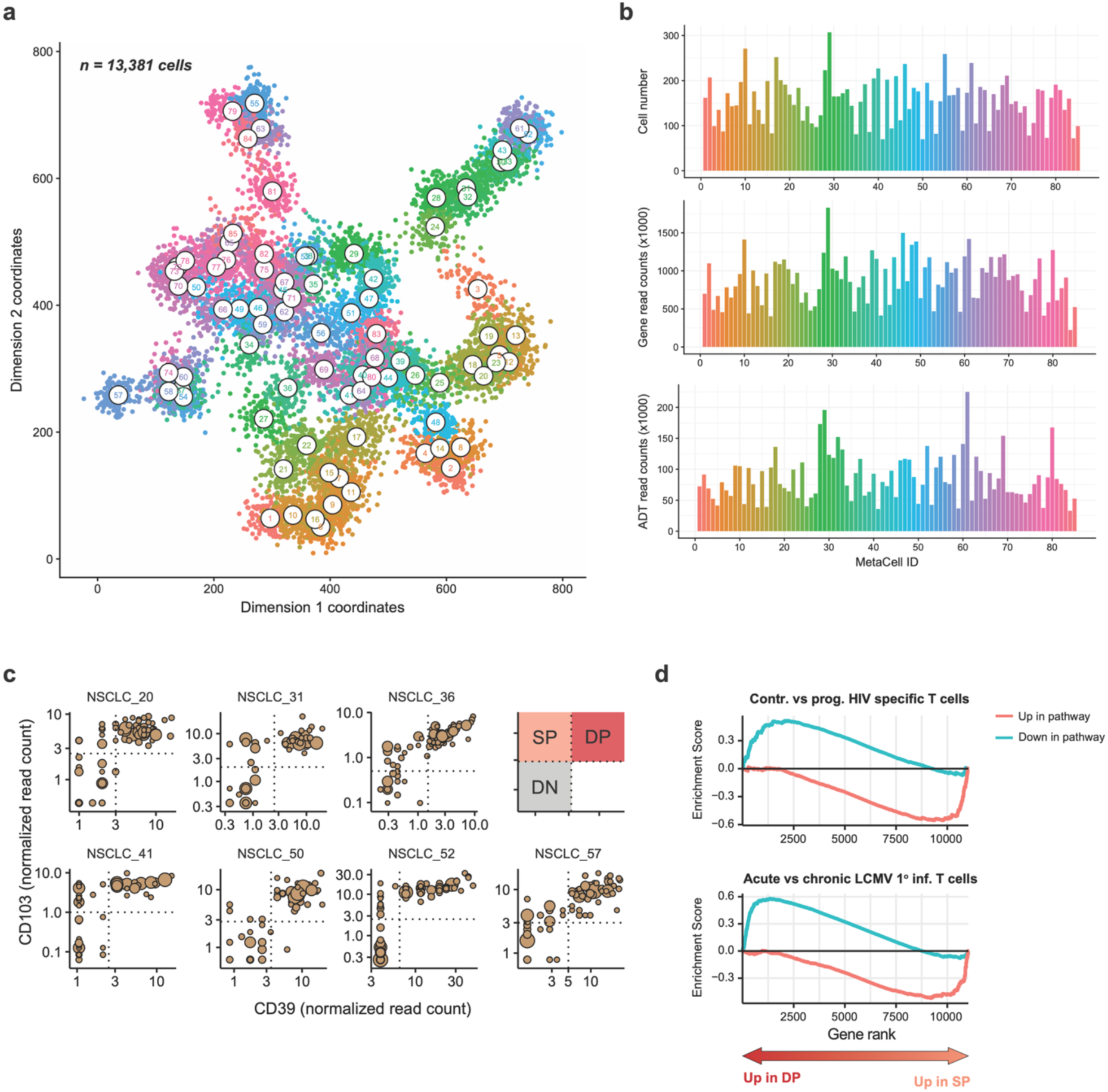
Single-cell RNA sequencing of tumor derived. CD8^+^ **T cells from non-small cell lung cancer lesions (NSCLC).** 10X single-cell RNA sequencing (scRNAseq) coupled with TCR profiling and antibody-derived tags (ADT) was performed on CD8^+^ T cells derived from non-small cell lung cancer lesions. (**a**) 2D projection of the transcriptome data of all measured single cells. Cells are colored by corresponding MetaCell. (**b**) Total cell numbers, transcriptome read counts, and ADT read counts per MetaCell. (**c**) ADT count thresholds under to define the CD103^NEG^CD39^NEG^ (DN), CD103^POS^CD39^NEG^ (SP), and CD103^POS^CD39^POS^ (DP) **CD8^+^ T cell** populations per patient. (**d**) Gene-set enrichment analysis for the MSigDB (IMMUNESIGDB) gene-sets ‘CONTROLLER VS PROGRESSOR HIV SPECIFIC CD8 TCELL’ and ‘ACUTE VS CHRONIC LCMV PRIMARY INF CD8 TCELL’. Color lines indicate the ‘UP’ and ‘DOWN’ versions of the gene-sets.

**Supplementary Figure 2.**
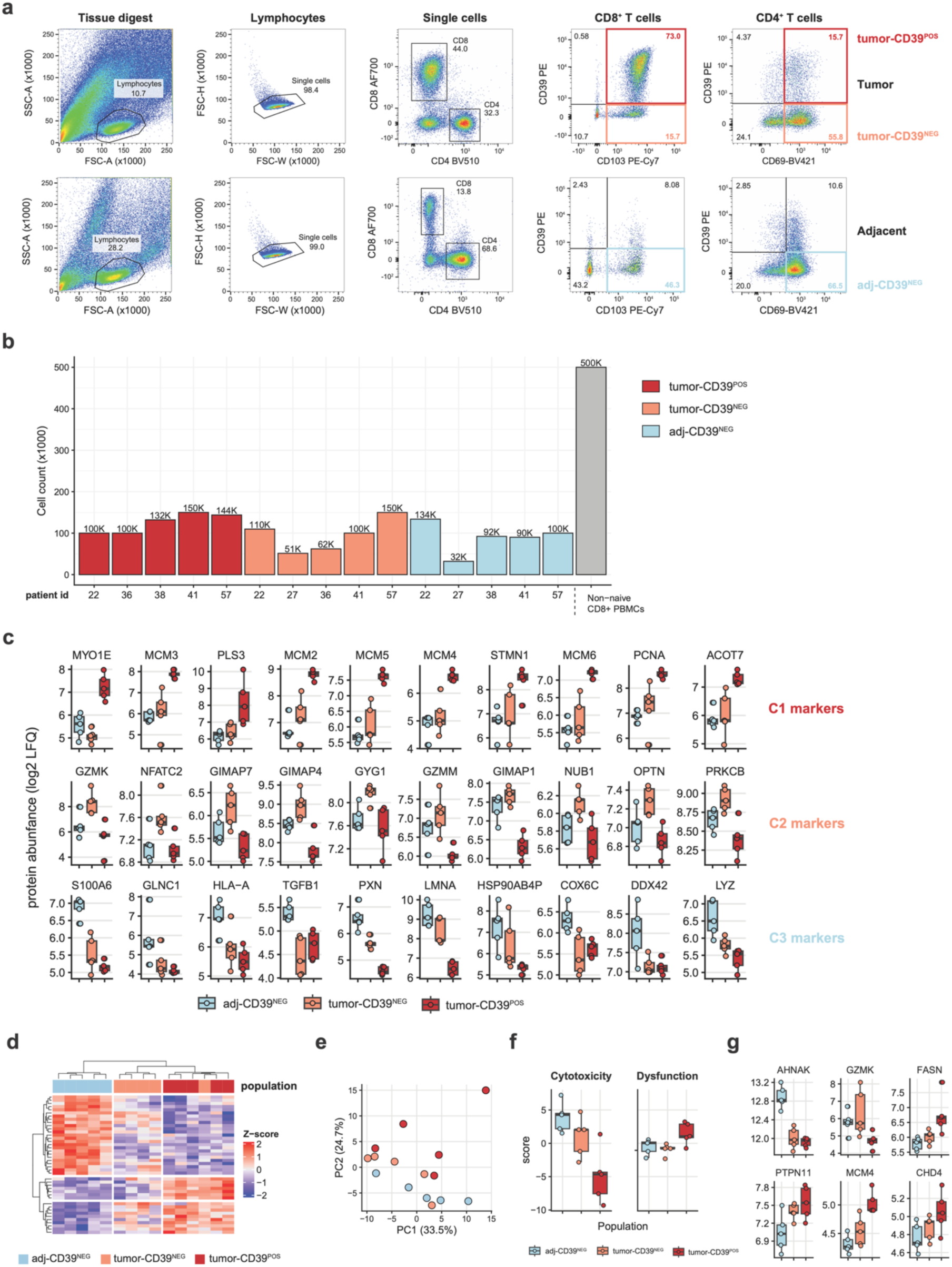
Proteomics of cancer-associated T cell populations. (**a**) Flow-based sorting strategy. Purified populations are indicated in colored boxes. (**b**) Cell numbers obtained per population. (**c**) Top 10 marker proteins (based on fold-difference) for each protein expression cluster (Figure 1j). Dots indicate populations from individual patients (n = 5). (**d-g**) TMT-16 LC-MS performed on “bystander” CD69⁺CD39⁻ single-positive (tumor-CD39^NEG^) T cells, “dysfunctional” CD69⁺CD39⁺ double-positive (tumor-CD39^POS^), and “tissue-patrolling” CD69⁺CD39⁻ (adj-CD39^NEG^) CD4^+^ cancer-associated T cells. (**d**) Hierarchical clustering of all differentially expressed proteins across CD4^+^ T cell populations indicated in (a). (**e**) Principal component analysis of the top 500 variable proteins. Dots indicate individual samples. (**f**) Dysfunction and cytotoxicity scores (summed z-scores across genes within gene-set) for indicated CD4^+^ T cell populations. (**g**) Selected proteins involved in T cell function that displayed significant differential expression across the assessed populations. Boxplots (**c, f, g**) show the median and 25th/75th percentiles, whiskers extend to 1.5× the interquartile range.

**Supplementary Figure 3.**
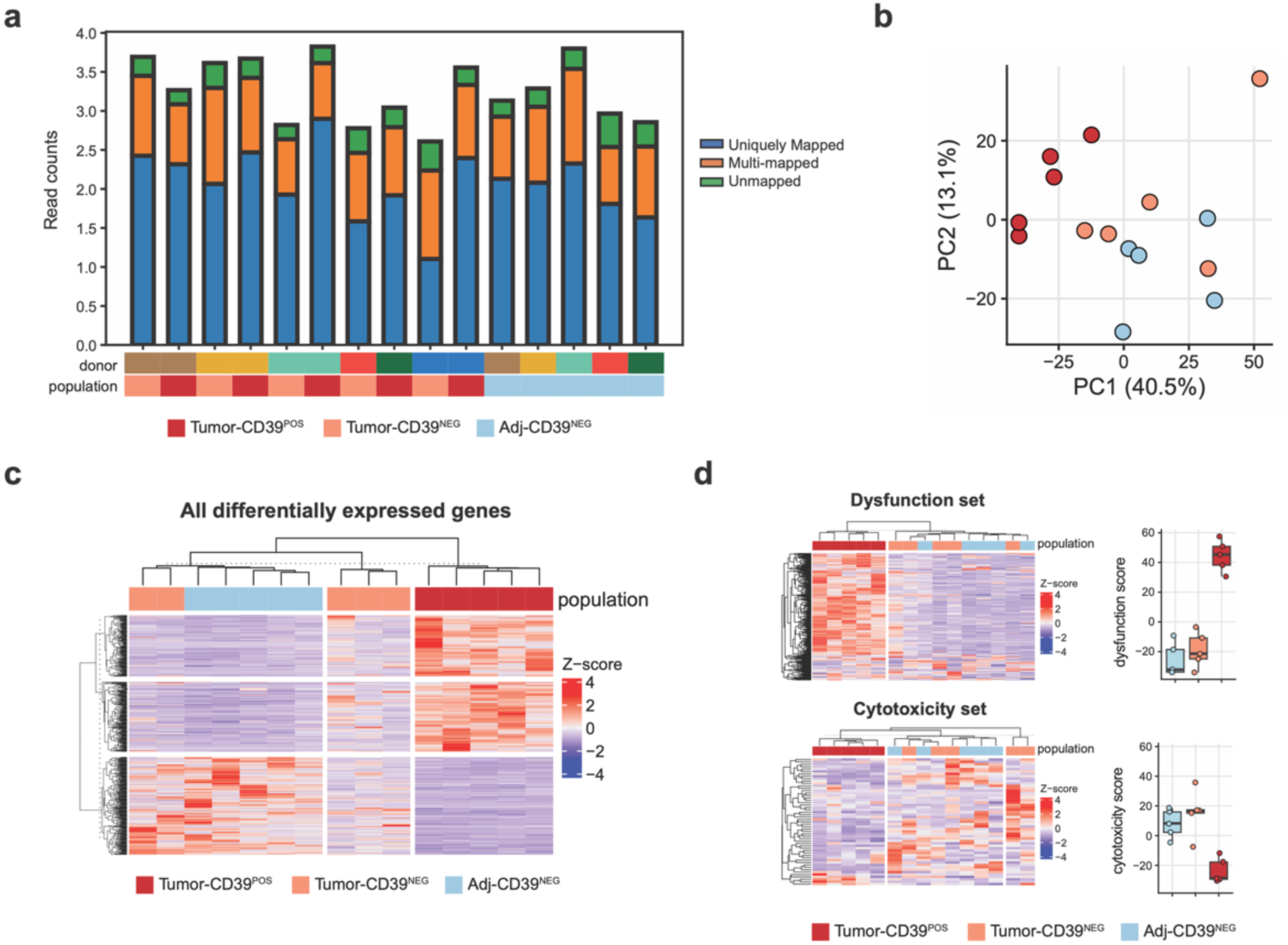
Bulk RNA sequencing of cancer-associated CD8^+^ T cell populations. Bulk RNA sequencing (SMARTseq2 protocol) was performed on indicated CD8^+^ T cell populations from NSCLC lesions. (**a**) Read alignment of indicated samples. (**b**) Principal component analysis of the top 2,000 variable transcripts. Dots indicate individual samples. (**c**) Hierarchical clustering of all differentially expressed proteins across the assessed populations. (**d**) Hierarchical clustering of proteins associated with T cell cytotoxicity and dysfunction (left). Dysfunction and cytotoxicity scores for the indicated populations (right). Scores were calculated per patient as the summed z-scores of genes within the respective dysfunction or cytotoxicity gene sets. Boxplots show the median and 25th/75th percentiles, whiskers extend to 1.5× the interquartile range.

**Supplementary Figure 4.**
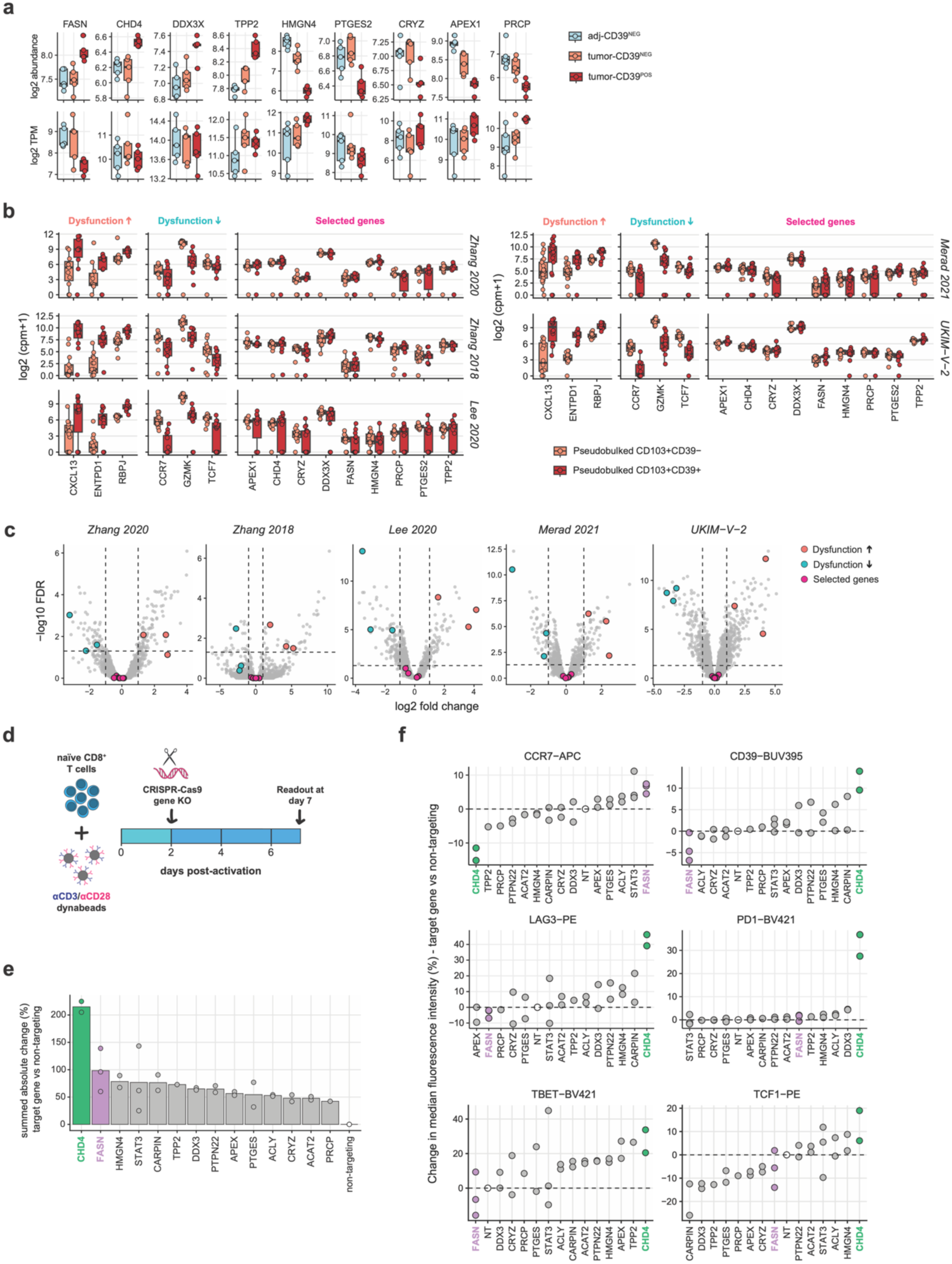
Genetic perturbation of FASN and CHD4 influences T cell state. (**a**) Proteins selected for functional follow-up experiments. Protein (top) and mRNA (bottom) abundance are shown. Dots indicate populations from individual patients. (**b-c**) Re-analysis of scRNAseq data from 5 studies included in the meta-analysis of Salcher et al. (Salcher et al., 2022, Cancer Cell, 10.1016/j.ccell.2022.10.008). CD103^NEG^CD39^NEG^ and CD103^NEG^CD39^POS^ CD8^+^ T cell populations were defined based on mRNA expression, followed by pseudo-bulking (b) and differential expression analysis (c). Selected genes for follow-up, as well as genes positively and inversely associated with dysfunction, are indicated in the plots. In boxplots, dots indicate T cell populations from individual patients. (**d-f**) Human naïve CD8⁺ T cells were activated with anti-CD3/CD28 Dynabeads and nucleofected at day 2 post-activation with either non-targeting Cas9–guide RNA complexes or complexes targeting the indicated genes. Expression of six surface proteins and five transcription factors was measured by flow cytometry at day 7 post-activation. (**d**) Experimental timeline. (**e**) Absolute percentage change in marker expression (median fluorescence intensity) relative to non-targeting control treated CD8^+^ T cells, averaged across all assessed proteins. Dots represent individual donors (n = 2-3). (**f**) Percentage change in protein expression (median fluorescence intensity) relative to non-targeting control. Dots represent individual donors (n = 2-3). Boxplots (**a, b**) show the median and 25th/75th percentiles, whiskers extend to 1.5× the interquartile range. P values were determined using edgeR’s quasi-likelihood F-test framework using the Benjamini–Hochberg method for multiple testing correction (**c**).

**Supplementary Figure 5.**
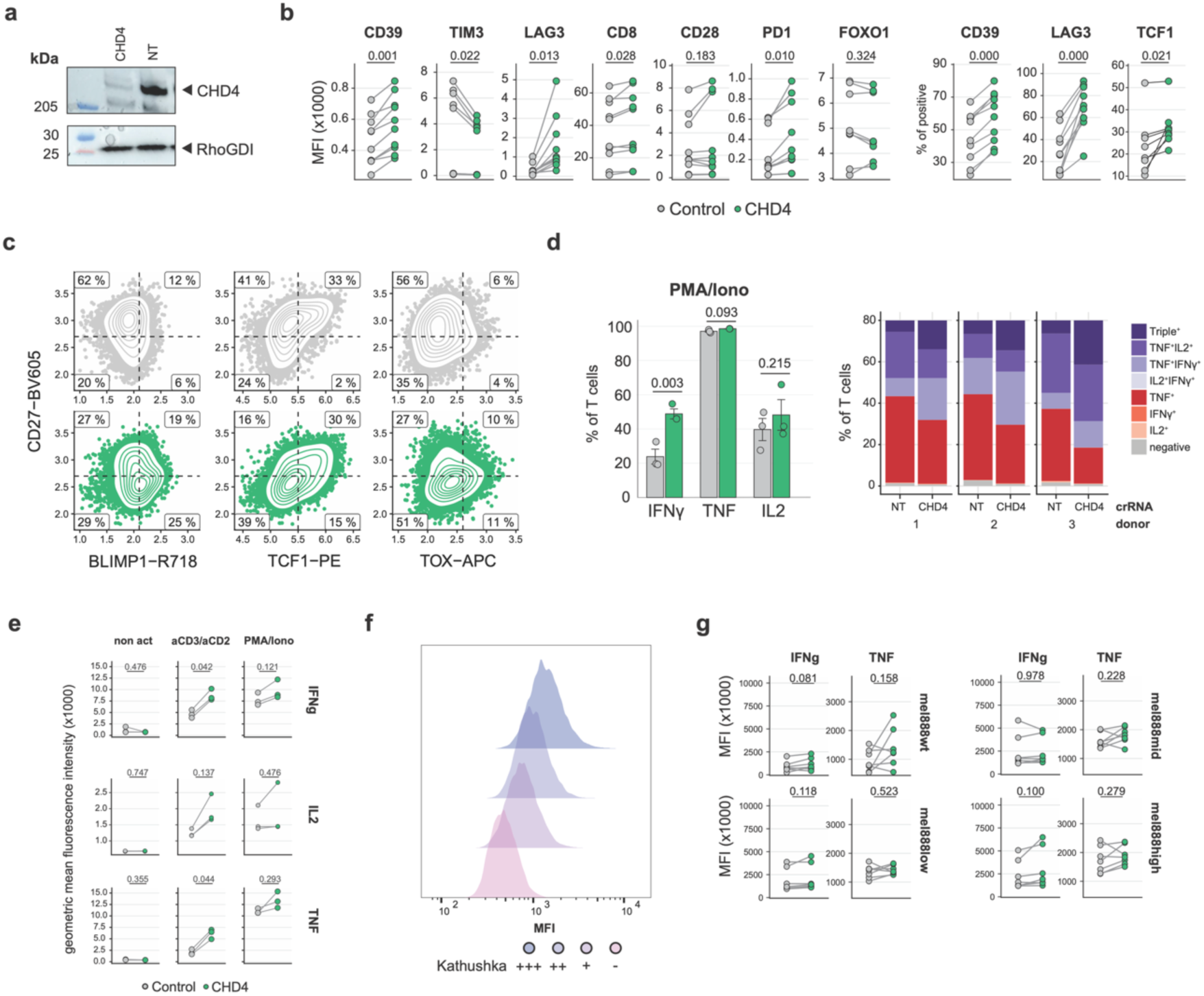
CHD4 regulates T cell effector responses upon antigen recognition. (**a**) Immunoblot confirming CHD4 protein loss in edited cells relative to control. RhoGDI serves as a loading control. (**b-d**) Protein expression levels for indicated surface proteins and transcription factors (**b-c**) and cytokines (**d**), measured by flow cytometry. Summarizing strip charts (b, d) and density plots (c) are shown. Lines connect samples obtained from individual donors (n = 3-10). (**e**) Representative density plots showing fluorescence intensity (MFI) of kathuska in Mel888 cells that were engineered to express a membrane-tethered single-chain anti-CD3 antibody at different expression levels. (**f**) Intracellular cytokine staining of indicated cytokines after a 16-hour co-culture (E:T of 1:1) with engineered Mel888 cells. Lines connect samples obtained from individual donors (n = 7). Displayed graphs are representative of 2-4 independent experiments, and contain data from 1 experiment (**d, e**) or are compiled from 3 (**b, c, f**) experiments. P values (**b, d, e**) were determined by multiple paired two-sided Student’s t-tests with Benjamini–Hochberg FDR correction for multiple testing.

**Supplementary Figure 6.**
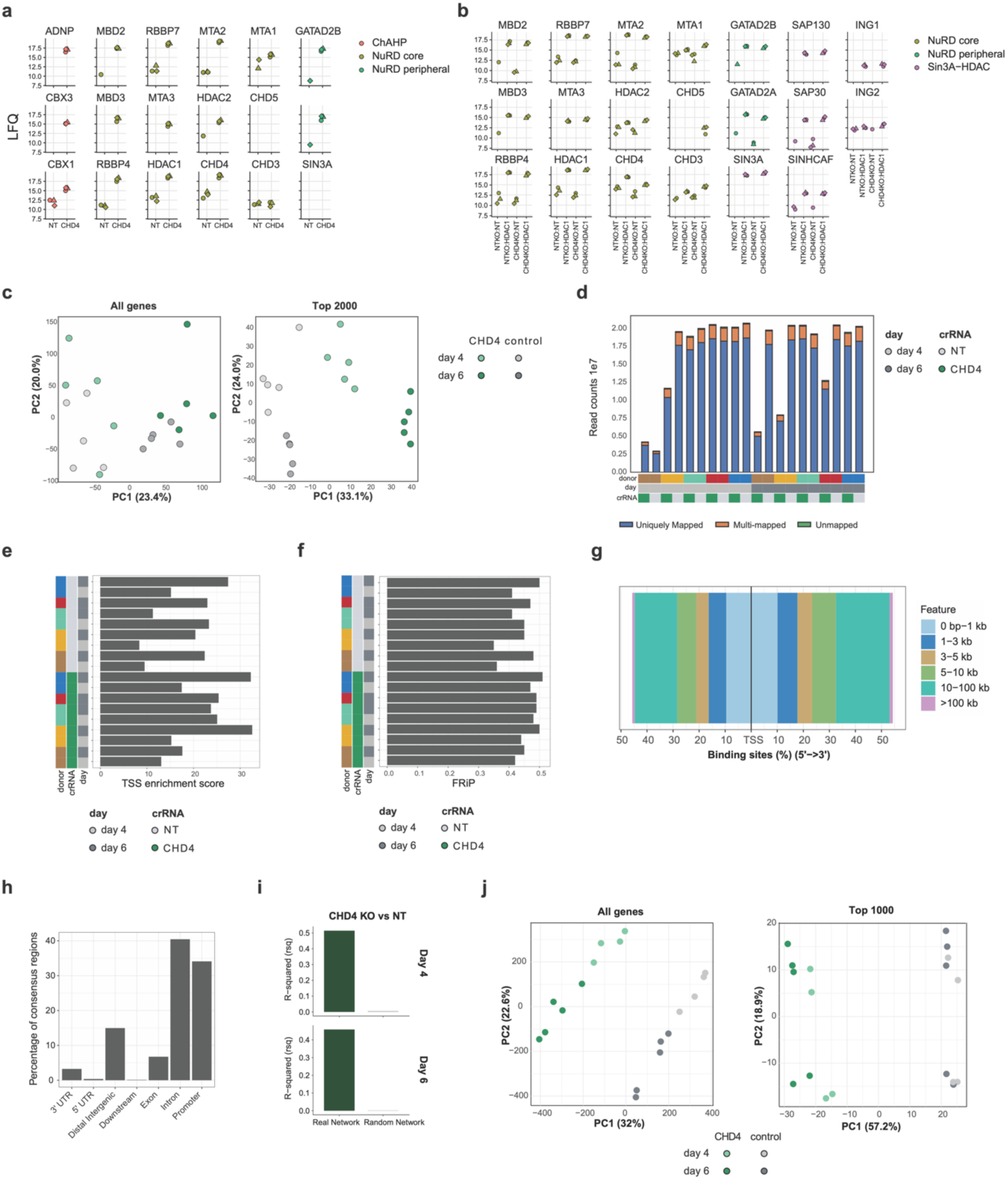
CHD4-dependent control of NuRD complex function and epigenomic remodeling in T cells. (**a**) Co-immunoprecipitation on T cell lysates (day 7 post activation) using an anti-CHD4 antibody and isotype control, followed by mass spectrometry–based identification of interacting proteins, highlighting NuRD complex components (n = 3). (**b**) Co-immunoprecipitation on T cell lysates using an anti-HDAC1 antibody and isotype control on CHD4-proficient and -deficient T cells (n = 3). (**c**) RNA-seq and ATAC-seq was performed on CHD4-proficient and -deficient T cells (from matched donors) at day 4 and day 6 post activation, and principal component analysis (PCA) of the RNA-seq data was used to assess global transcriptional variation using all and top 2000 most variable genes. Points indicate samples obtained from individual donors (n = 4-5). (**d**) Sequencing quality control metrics for RNA-seq libraries, showing read alignment statistics (uniquely mapped, multi-mapped and unmapped reads). (**e**) TSS enrichment scores for ATAC-seq samples. (**f**) Fraction of reads in peaks (FRiP) for ATAC-seq samples. (**g**) Differential accessibility of regions identified by ATAC-seq, shown as distribution of binding sites relative to transcription start sites (TSS). (**h**) Genomic annotation of identified accessible consensus regions. (**i**) Bar plots show enhancer-based gene regulatory network (eGRN) performance evaluated by R² prediction accuracy, comparing the real network to randomized networks at day 4 and day 6. (**j**) PCA of ATAC-seq chromatin accessibility profiles based on the top 1000 most variable peaks across samples. Points indicate samples obtained from individual donors.

**Supplementary Figure 7.**
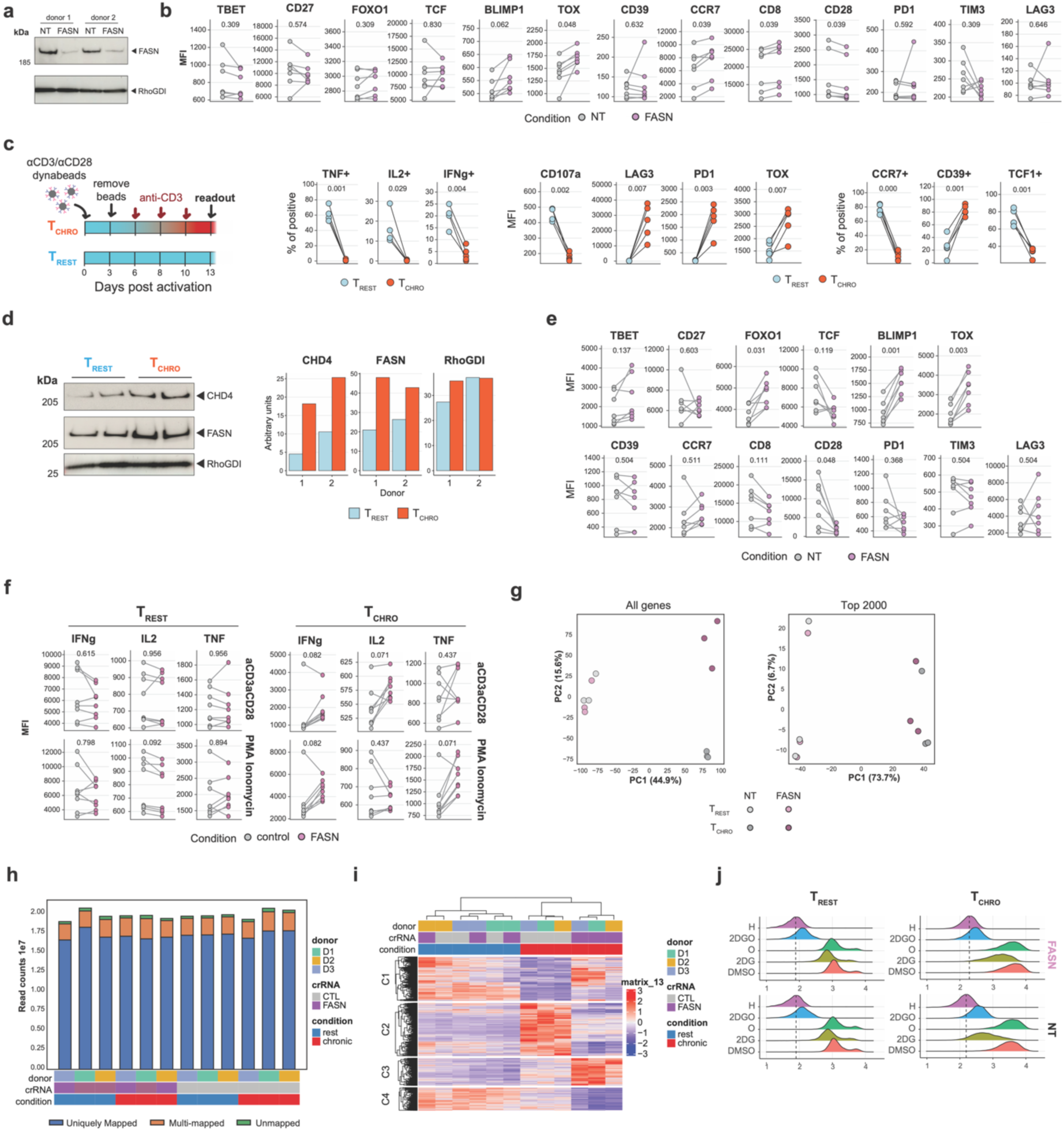
FASN-dependent control of CD8^+^ T cell function and metabolism. (**a**) Immunoblot confirming FASN protein loss in edited CD8^+^ T cells relative to control. RhoGDI serves as loading control. (**b**) Relative protein expression levels for indicated transcription factors and surface markers of resting (T_REST_) cells measured by flow cytometry. Paired dots represent individual donors under FASN-proficient or -deficient conditions (n = 7). (**c**) Activated CD8^+^ T cells were cultured in the presence or absence of plate-bound anti-CD3 antibodies for 7 days and analyzed at day 14 post-activation. Quantification of cytokine production (TNF, IL2, IFNγ) and degranulation (CD107a) alongside phenotypic markers of T_REST_ and chronically activated (T_CHRO_) cells by flow cytometry. Lines connect matched donor samples across conditions (n = 5). (**d**) Western-blot quantification of indicated proteins in total cell lysates of T_REST_ and T_CHRO_ cells. Representative western-blot (left) and quantified intensity (right) are shown. (**e**) Relative protein expression levels of indicated transcription factors and surface markers of T_CHRO_ cells, assessed by flow cytometry. Paired donor samples are shown for FASN-proficient and - deficient cells (n = 7). (**f**) Intracellular cytokine staining following 3-hour restimulation with anti-CD3/CD28 antibodies or PMA/ionomycin in the presence of brefeldin A, comparing T_REST_ and T_CHRO_ cells. Dots indicate individual donors, with paired comparisons between FASN-proficient and -deficient conditions. (**g**) Principal component analysis (PCA) of RNA-seq datasets from T_REST_ and T_CHRO_ cells, shown for all genes (left) and the top 2000 variable genes (right). Each point represents an individual donor sample (n = 3). (**h**) RNA-seq mapping statistics showing proportions of uniquely mapped, multi-mapped, and unmapped reads across samples. (**i**) Heatmap of differentially expressed genes across donors and conditions with hierarchical clustering. (**j**) SCENITH analysis of metabolic dependencies in T_REST_ and T_CHRO_ CD8^+^ T cells. Cells were treated for 15 minutes with metabolic inhibitors targeting glycolysis (2-deoxyglucose; 2DG), mitochondrial respiration (oligomycin; O), both pathways (2DGO), translation (harringtonine; H), or left untreated (DMSO), followed by puromycin incorporation for 20 minutes to assess protein synthesis rates by flow cytometry. Density plots show puromycin staining under each condition for FASN-KO and NT control T cells. Displayed graphs are representative of 2-4 independent experiments, contain data from 2 (**c**) or 3 (**b, e, f**) experiments, or represent a single experiment (**g, h, i**). P values (**b, c, e, f**) were determined by multiple paired two-sided Student’s T tests with Benjamini–Hochberg FDR correction for multiple testing.

## Supplementary Data 1

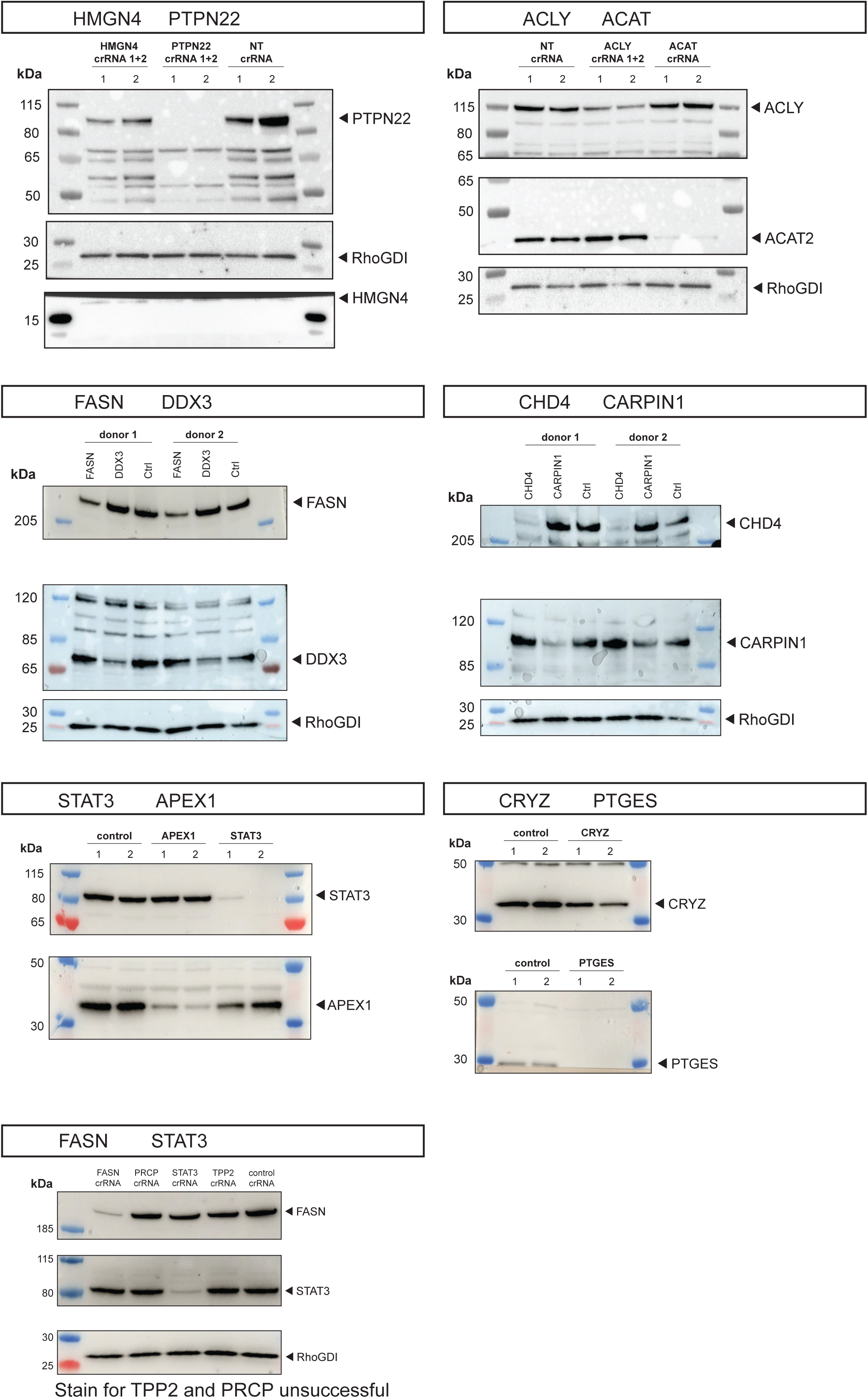

## Supplementary Table descriptions

|  |  |
| --- | --- |
| Dys_cytox_scores.xlsx | Supplementary Table 1 |
| proteomics_NSCLC_Tcell_abundance_CD8.tsv | Supplementary Table 2 |
| proteomics pathway analysis | Supplementary Table 3 |
| proteomics_NSCLC_Tcell_populations_CD4.tsv | Supplementary Table 4 |
| DE of all proteomics and RNAseq | Supplementary Table 5 |
| NSCLC_Proteomics_pathways.tsv | Supplementary Table 6 |
| MFI_combined_surface_tf.csv | Supplementary Table 7 |
| Pulldown_CHD4_abundance.tsv | Supplementary Table 8 |
| Pulldown_HDAC1_abundance.tsv | Supplementary Table 9 |
| CHD4 DE genes | Supplementary Table 10 |
| CHD4 DAR | Supplementary Table 11 |
| CHD4 pathways | Supplementary Table 12 |
| FASN DE genes | Supplementary Table 13 |
| FASN pathways | Supplementary Table 14 |
| Antibodies | Supplementary Table 15 |
| CRISPR guides | Supplementary Table 16 |

